# Targeted metagenomic sequencing with spiked primers for enrichment of viruses in wastewater for pathogen surveillance

**DOI:** 10.64898/2025.12.26.690930

**Authors:** Matt C. Parker, Nisachon Apinda, Siriwat Uamngamsup, Polawat Phetra, Terdsak Yano, Kanokwan Sangkakam, Thanaporn Eiamsam-ang, Anucha Muenthaisong, Pune Phetra, Angkana T. Huang, Mark Smolinski, Sarah H. Olson, Nathan D. Grubaugh, Maira Phelps, Karyna Rosario, Juliana Gil, Patipat Susumpow

## Abstract

**Background:** Wastewater surveillance offers an underutilized opportunity to identify high-risk viral pathogens that pose public health risks. Although metagenomic approaches have been increasingly adopted for human wastewater surveillance, little attention has been given to its application in rural agricultural settings. Untargeted metagenomic sequencing of wastewater poses considerable technical challenges for viral detection due to fragmented genomes and low viral abundance. While existing enrichment methods partially address these challenges, high costs and proprietary protocols limit adoption in resource-constrained settings.

**Objective:** We developed a fully open-source primer design algorithm (*Open MSSPE Design*) and evaluated Metagenomic Sequencing with Spiked Primer Enrichment (MSSPE)^1^, as a practical strategy for metagenomic surveillance of swine slurry and farm wastewater samples collected from rural agricultural settings. This non-invasive MSSPE strategy addresses the challenge of detecting extremely low-abundance viral targets amid samples dominated by a background of bacterial and eukaryotic nucleic acids.

**Methodology:** We generated fifteen primer sets targeting high-priority DNA and RNA viral pathogens affecting human and animal health in Southeast Asia using our open-source primer design algorithm. Twenty-five wastewater and swine slurry samples from smallholder farms in northern Thailand underwent parallel library preparation—untargeted mNGS and MSSPE—for direct comparison. Libraries were sequenced on Illumina platforms and analyzed using Chan Zuckerberg ID (CZ ID)^2^. Rarefaction analysis assessed performance at sequencing depths of 100,000–1.5 million reads per sample.

**Results:** We detected multiple high-risk DNA and RNA viruses in wastewater samples from smallholder farm operations. MSSPE achieved substantial viral enrichment across nine pathogenic DNA and RNA viruses, with a median two-fold enrichment of reads per million, with variability across targets (IQR: 1.01–3.44x) and a nearly 10% median increase in breadth of genome coverage (IQR: 4.54–11.84%), while retaining sensitivity for untargeted pathogens. MSSPE also increased the odds of detecting targeted viruses (OR 1.35, CI 1.14–1.60), with the greatest advantage at shallow sequencing depths where MSSPE required fewer reads to identify targeted viral taxa relative to mNGS.

**Conclusions:** MSSPE demonstrated the ability to enrich shallow-depth sequencing (<2M reads per sample) sufficiently to detect priority viruses without substantially increasing library preparation time or cost. This open-source workflow supports cost-effective metagenomic viral surveillance for resource-constrained settings, providing a non-invasive method for detecting low-abundance viral targets in high-background sample types at rural agricultural interfaces with elevated risk of zoonotic spillover.

## 1. INTRODUCTION

Animal disease outbreaks are a growing threat to global food security, biosecurity, and public health^3,4^. Most livestock in low- and middle-income countries (LMICs) are raised in backyard farming systems, with minimal biosecurity and frequent contact between domestic animals, wildlife, and nearby human communities^5^. Such conditions increase the likelihood of cross-species spillover^21^ and disproportionately impact smallholder farmers, who experience more frequent outbreaks and face greater financial barriers to recovery^6^. Thailand exemplifies this risk landscape: human cases of highly pathogenic avian influenza A (H5N1) have been reported in association with poultry^7^, and *Henipavirus nipahense* (Nipah virus) has been detected in multiple Thai bat populations^8^, many of which overlap geographically with swine operations—a known amplifier of Nipah transmission^9^. Despite this clear risk at a critical animal–human interface, systematic surveillance in small-scale agricultural settings remains limited.

Wastewater surveillance has emerged as a powerful tool in human health systems^10^, consistently providing critical lead time ahead of clinical case surges^11,12^, and enabling proactive public health responses independent of case-based surveillance. Evidence suggests that sparse, non-consecutive wastewater sampling can still reliably detect changes in viral abundance, reducing resource demands and improving feasibility in resource-limited settings^13^. However, the epidemiological granularity of conventional wastewater surveillance systems often encompasses catchment areas that are too large to support direct, targeted interventions.

In targeted application, wastewater can be leveraged as a non-invasive alternative to individual diagnostic testing^14^, offering a narrower focus for immediate, actionable responses directly at the source of potential outbreaks. Controlling small-scale outbreaks is an effective strategy for preventing large-scale epidemics and pandemics, as early intervention at localized transmission interfaces can prevent epidemic amplification^15^. A farm-specific surveillance approach aligns closely with this principle, providing the focused catchment size necessary for rapid, direct intervention when pathogenic threats are detected, rather than the broad population-level monitoring that characterizes traditional wastewater surveillance systems. One of the most promising approaches in our biosecurity arsenal to detect and mitigate future outbreaks is the application of metagenomics in these settings, given its capability to identify both emerging and novel pathogens. When applied non-invasively via wastewater in high-risk environments, such as livestock farms or live animal markets—where animals are most likely to come into contact with humans—such a system could help interrupt transmission pathways at a critical interface.

Untargeted and targeted metagenomic sequencing of wastewater has shown promise for pathogen surveillance, but the real-world application has been limited^16,17^. Untargeted metagenomic next-generation sequencing (hereafter mNGS) has been shown to require substantial sequencing depth to detect pathogenic viruses in wastewater, resulting in high per-sample costs. Wastewater mNGS samples are often dominated by bacteria (>99%)^16^ and viral concentrations are extremely low, with the majority of viral reads consisting of plant viruses and bacteriophages^18^ rather than pathogenic viruses of concern. Targeted metagenomics can potentially reduce the need for expensive deep sequencing to capture viral genomes in low abundance, significantly reducing cost barriers to utility. However, current targeted approaches present distinct trade-offs between sensitivity, specificity, and practical implementation^19^. Amplicon-based methods, while capable of generating comprehensive genome coverage for known pathogens, exhibit limited capacity to detect highly divergent genomes, potentially missing novel species with pandemic potential^20^. Alternatively, hybrid capture probe systems can accommodate greater genomic diversity but remain cost-prohibitive for most resource-constrained laboratories and introduce additional complexity and processing time to surveillance workflows^21^.

The metagenomic sequencing with spiked primer enrichment (MSSPE) method may provide an alternative to mNGS, viral capture, or even multiplex PCR offering high viral yield at low sequencing depths without substantially increasing overall costs. MSSPE preferentially enriches multiple viral genomes by spiking primers targeting overlapping regions across viral genomes of interest alongside random hexamers typically used in mNGS protocols. The spiked primers function as reverse-transcription priming oligos and PCR primers, thus enriching specific targets, while random hexamers can recover untargeted genomes. Because MSSPE requires only a pool of synthesized primers (estimated cost of US$125 per pool) in addition to mNGS reagents, MSSPE is potentially well suited for wastewater samples from small-scale livestock farms, where sequencing resources are limited. However, application of MSSPE has so far been limited to low-complexity sample types from high-income clinical settings^1^ and proprietary methods have made reproducing such MSSPE protocols a barrier to adoption^22^.

Here, we present the development and preliminary validation of an open-source MSSPE approach, through a pilot study monitoring viral pathogens in small, rural farms located in northern Thailand. We used the assay to enrich 15 human and animal viral pathogens from swine slurry and farm wastewater samples. By integrating open-source primer design with a low-cost, field-adaptable workflow, we demonstrate that MSSPE can enhance detection of priority viral pathogens in low-abundance, high-background samples from rural agricultural settings.

## 2. RESULTS

### 2.1 Open MSSPE Design as an open-source pipeline for targeting viral pathogens

We developed a new primer design algorithm and packaged it as an open-source, flexible, and automated software called *Open MSSPE Design* to obtain primers optimized for MSSPE (**Fig. 6**). We used *Open MSSPE Design* to generate fifteen primer sets, each targeting a priority RNA and DNA viral genome (**Table 1**). We selected priority pathogens based on criteria derived from the most recent multi-sectoral One Health Zoonotic Disease Prioritization (OHZDP) workshop for Thailand^23^, as well as priority pathogen families identified by the WHO’s R&D Blueprint for Epidemics^24^. In addition, we selected viruses representing diverse genomic structures, such as segmented and non-segmented genomes, enveloped and non-enveloped viruses, and short (e.g. PCV ∼1 kb) and long (e.g. ASFV ∼190 kb) genomes. Ultimately, we selected viruses that were likely to be present on the farms based on a preliminary experiment, represented high-risk zoonoses endemic to Southeast Asia, or were highly pathogenic animal viruses with the potential to cause severe economic and public health harm in smallholder farming communities. We also included primers targeting the HPV18 oncogenes (E6 and E7) integrated in HeLa cells and tested enrichment using HeLa cell RNA to assess method performance. HPV18 is genetically dissimilar to other targeted pathogens and was unlikely to be present in swine farms in Thailand.

**Table 1.** List of primer sets designed for 15 targeted viral pathogens used in MSSPE library preparation. All primers were 15 nucleotides long. Viral taxa with asterisk(*) denote taxa which were targeted due to the likelihood of presence on farms based on preliminary experiments. HPV18 primers were designed to target the E6 and E7 genes of the viral genome. IAV primers targeted all non-subtyping segments (1-3,5,7,8) as well as segments (4,6) for H1, H5, and H7 subtypes. All other primers were designed using whole genome references. *Primers per kb* genome shows the number of primers per mean length of reference genome, divided by 1000. *Refs per kb genome* shows the number of references per mean length of reference genome, divided by 1000.

| Virus | Total Primers | Forward | Reverse | References | Primers / kb genome | Refs / kb genome |
| --- | --- | --- | --- | --- | --- | --- |
| African swine fever virus (ASFV) | 212 | 106 | 106 | 122 | 1.18 | 0.68 |
| Classical swine fever virus (CSFV) | 109 | 47 | 62 | 219 | 8.86 | 17.80 |
| E6/E7 oncogenes of HeLa cells (HPV18) | 5 | 1 | 4 | 2 | 6.29 | 2.52 |
| Enterovirus G (EV-G) | 116 | 56 | 60 | 44 | 15.77 | 5.98 |
| Foot-and-mouth disease virus (FMDV) | 48 | 22 | 26 | 72 | 5.78 | 8.67 |
| Influenza A virus (IAV) | 631 | 292 | 339 | 3,613 | 46.74 | 267.63 |
| Nipah virus (NiV) | 111 | 61 | 50 | 80 | 6.17 | 4.44 |
| Porcine astrovirus (PAstV) | 52 | 21 | 31 | 16 | 7.99 | 2.46 |
| Porcine bocavirus (PBoV) | 37 | 20 | 17 | 16 | 7.12 | 3.08 |
| Porcine circovirus (PCV) | 17 | 10 | 7 | 191 | 9.62 | 108.09 |
| Porcine kobuvirus (PKV) | 61 | 24 | 37 | 64 | 7.53 | 7.90 |
| Porcine reproductive and respiratory syndrome virus (PRRSV) | 64 | 32 | 32 | 118 | 4.18 | 7.71 |
| Porcine sapelovirus (PSapeV) | 116 | 53 | 63 | 40 | 15.43 | 5.32 |
| Porcine sapovirus (PSapoV) | 50 | 32 | 18 | 21 | 6.70 | 2.82 |
| Porcine teschovirus (PTeV) | 128 | 52 | 76 | 27 | 17.76 | 3.75 |

We concentrated samples using a simplified polyethylene glycol (PEG) precipitation method, and conducted paired experiments in which two laboratory staff worked in parallel on each sample to compare enrichment performance between MSSPE and mNGS. MSSPE libraries were spiked with the designed primers during library preparation. Taxa were then identified using the cloud-based taxonomic classification platform Chan Zuckerberg ID (CZ ID) (**Fig. 1**).

**Figure 1.**
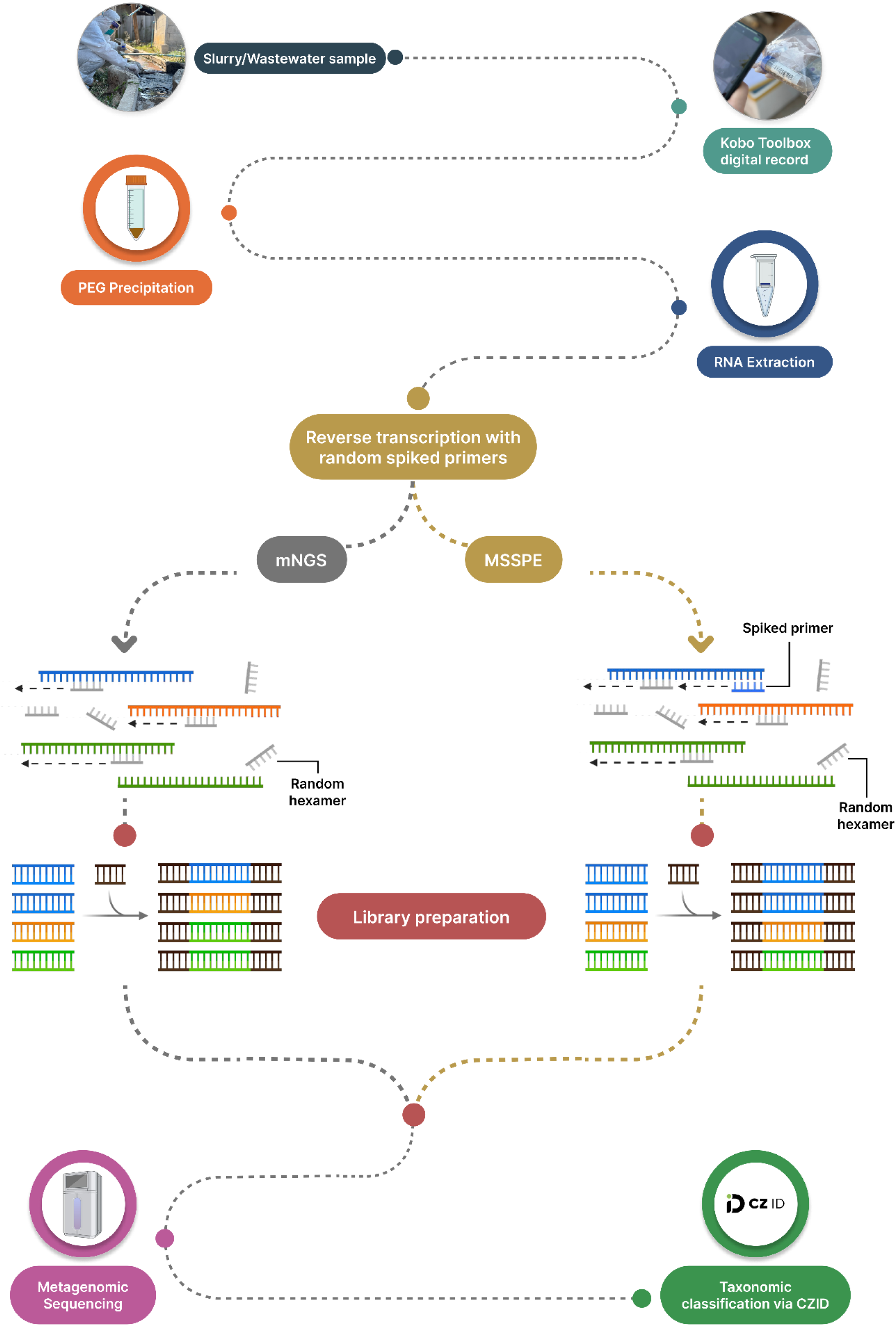
Complete experimental workflow: sample collection, mobile field data collection, PEG precipitation, RNA extraction, paired library preparations (mNGS and MSSPE), metagenomic sequencing, and taxonomic classification analysis via CZ ID.

### 2.2 MSSPE enriched viral reads compared to mNGS

We detected nine of the fifteen viral pathogens targeted via spiked primers: Porcine Teschovirus (TeV), Porcine Kobuvirus (PKV), Porcine Sapporo Virus (PSapoV), Porcine Sapelovirus (PSapeV), Porcine Astrovirus (PAstV), Porcine Bocavirus (PBoV), Enterovirus G (EV-G), Porcine Circovirus (PCV), and HeLa cell DNA containing human papillomavirus 18 (HPV18) as a process control (**Fig. 2a, 2b; Table 2a, 2b**). Six targeted viral pathogens were not detected in either mNGS or MSSPE libraries: African swine fever virus (ASFV), Classical swine fever virus (CSFV), Foot-and-mouth-disease virus (FMDV), Nipah virus (NiV), Porcine reproductive and respiratory virus (PRRSV), and Influenza A virus (IAV).

**Figure 2.**
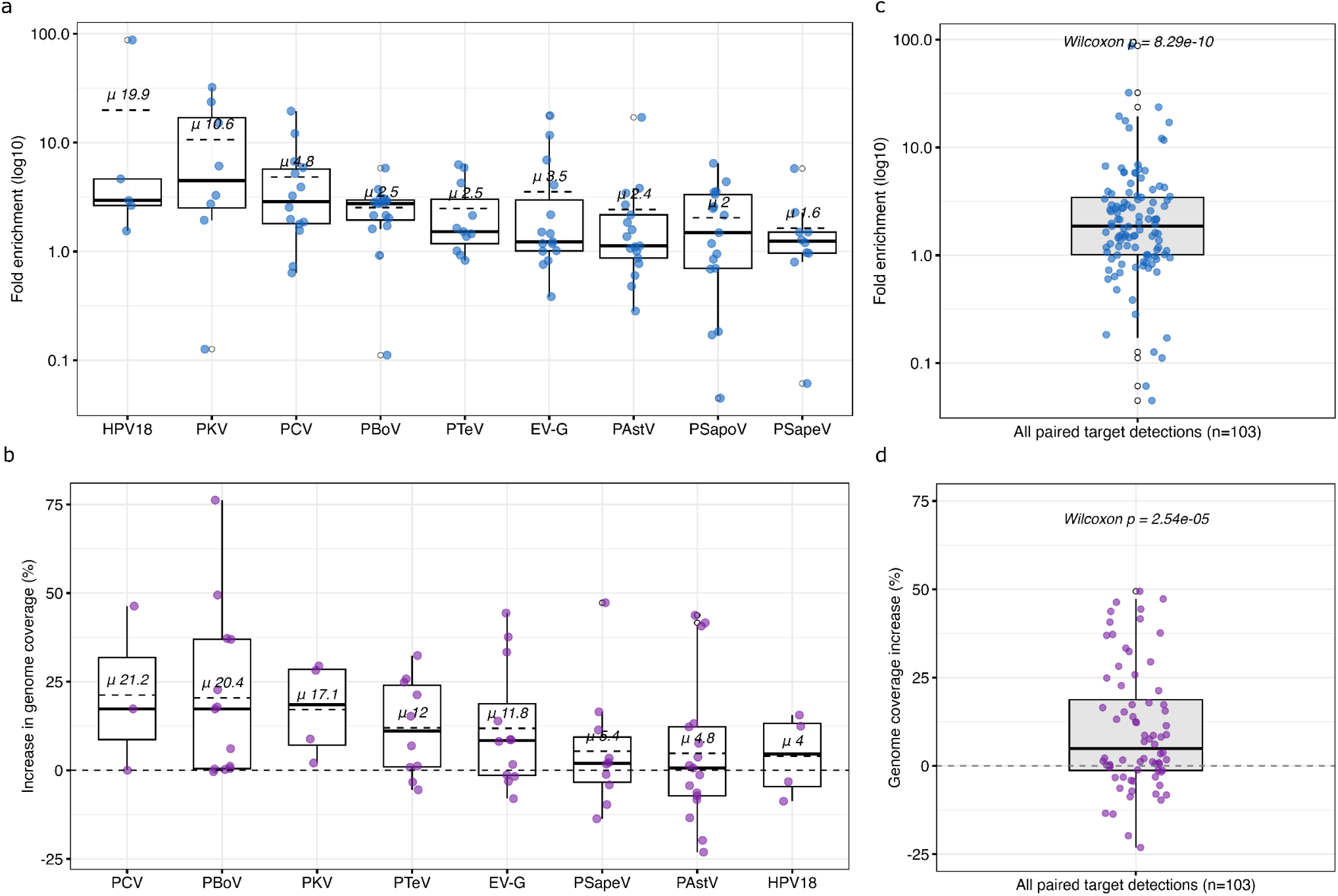
**(a)** Fold change (Log_10_ scale) in median reads per million (rPM) aligned to target viruses with spiked primers (MSSPE). Each point represents a paired sample’s fold change in rPM for a specific virus. Mean fold change is shown in dashed line. **(b)** Genome coverage breadth increase per viral target. Mean increase is shown above the median line.

**Table 2.** **(a)** Enrichment results for each viral target by library prep method. For each paired sample, enrichment was quantified as the ratio of ERCC-normalized reads per million (rPM) obtained from the MSSPE library relative to the corresponding mNGS library. To account for zero-read cases, a small pseudocount (0.1 rPM) was added to both values.

| Virus | rPM (mNGS) | rPM (MSSPE) | Median Fold Change | Mean Fold Change | IQR | p-value | p-adjusted | effect size (r) |
| --- | --- | --- | --- | --- | --- | --- | --- | --- |
| PKV | 49.86 | 136.85 | 4.68 | 10.64 | 2.5–17.3 | 0.023 | 0.052 | 0.792 |
| HPV18 | 4.90 | 16.51 | 2.94 | 19.88 | 2.6–4.6 | 0.062 | 0.093 | 0.905 |
| PCV | 21.29 | 55.67 | 2.89 | 4.83 | 1.8–5.7 | 0.005 | 0.022 | 0.713 |
| PBoV | 197.96 | 460.66 | 2.76 | 2.53 | 1.9–3 | 0.000 | 0.000 | 0.840 |
| PTeV | 34.38 | 47.36 | 1.52 | 2.48 | 1.2–3.2 | 0.032 | 0.058 | 0.643 |
| PSapoV | 35.49 | 79.06 | 1.49 | 2.04 | 0.7–3.3 | 0.132 | 0.170 | 0.373 |
| PSapeV | 19.19 | 27.30 | 1.24 | 1.64 | 1–1.5 | 0.322 | 0.329 | 0.338 |
| EV-G | 77.92 | 228.82 | 1.23 | 3.54 | 1–3.1 | 0.015 | 0.045 | 0.616 |
| PAstV | 183.45 | 356.94 | 1.12 | 2.43 | 0.9–2.2 | 0.329 | 0.329 | 0.247 |

b
| Virus | % coverage (mNGS) | % coverage (MSSPE) | Increase (%) | IQR | p-value | p-adjusted | effect size (r) |
| --- | --- | --- | --- | --- | --- | --- | --- |
| PCV | 34.17 | 56.66 | 22.49 | 17.6–28.2 | 0.059 | 0.118 | 0.861 |
| PBoV | 56.90 | 74.51 | 17.61 | 0.4–29.8 | 0.006 | 0.048 | 0.719 |
| PKV | 11.44 | 20.60 | 9.16 | 2.3–9.4 | 0.019 | 0.067 | 0.757 |
| PTeV | 33.49 | 43.40 | 9.91 | 1.2–21.3 | 0.025 | 0.067 | 0.630 |
| EV-G | 40.96 | 48.62 | 7.66 | -3.1–9.9 | 0.205 | 0.328 | 0.323 |
| PSapeV | 32.81 | 36.37 | 3.56 | -3.8–3.3 | 0.616 | 0.663 | 0.143 |
| PAstV | 52.43 | 57.14 | 4.71 | -7–11.1 | 0.663 | 0.663 | 0.108 |
| HPV18 | 15.23 | 19.25 | 4.02 | -4.6–13.2 | 0.584 | 0.663 | 0.365 |

| Virus | MSSPE detections | mNGS detections | Odds Ratio (MSSPE/mNGS) | 95% CI Lower | 95% CI Upper |
| --- | --- | --- | --- | --- | --- |
| EV-G | 21 | 14 | 1.500 | 0.763 | 2.950 |
| HPV18 | 10 | 9 | 1.111 | 0.451 | 2.734 |
| PAstV | 108 | 86 | 1.256 | 0.946 | 1.667 |
| PBoV | 62 | 39 | 1.590 | 1.065 | 2.373 |
| PCV | 19 | 12 | 1.583 | 0.769 | 3.262 |
| PKV | 10 | 4 | 2.500 | 0.784 | 7.971 |
| PSapeV | 17 | 16 | 1.062 | 0.537 | 2.103 |
| PSapoV | 26 | 21 | 1.238 | 0.697 | 2.200 |
| PTeV | 37 | 29 | 1.276 | 0.785 | 2.075 |

Overall, our results show that MSSPE libraries had substantial enrichment across nine targeted DNA and RNA pathogenic viral species with a median rPM enrichment of 2x (IQR: 1.01–3.44x; Wilcoxon *p* <0.001) (**Fig. 2c**; **Table 2a**), and a median increase in genome coverage breadth by 10% (IQR: 4.54–11.84%; Wilcoxon *p* <0.001) (**Fig. 2d**; **Table 2b**). MSSPE libraries were also significantly more likely to detect the targeted viral species when compared to mNGS libraries (pooled OR 1.35, 95% CI 1.14–1.60; McNemar *p* <0.001) (**Table 2c**). Enrichment via MSSPE varied widely depending on the targeted viral species. MSSPE also showed a ∼5% mean increase in genome coverage depth, a 2.5-fold mean increase in nucleotide reads (NT) and a 2.1-fold mean increase in amino acid reads (NR) aligned to targeted viral taxa.

Paired sample percent increase values are shown as single points. **(c)** Overall enrichment across 9 targeted viruses with MSSPE. Fold change (Log_10_ scale) in reads per million (rPM) aligned to any of the targeted viral taxa. Wilcoxon p-value compares paired samples’ mean rPM values. Each point represents a paired sample’s fold change for a specific virus. **(d)** Overall increase in genome coverage breadth across all targeted viral taxa. Each point represents a paired sample’s percent change in genome coverage breadth for a specific virus. Wilcoxon p-value compares paired samples’ mean change in genome coverage breadth.

Porcine circovirus (PCV), a small, circular single-stranded DNA virus (*Circoviridae*) affecting livestock operations and causing gastrointestinal distress in pigs, and *postweaning multisystemic wasting syndrome* (PMWS) in piglets, was detected in 73% of samples, and showed strong enrichment in rPM (2.9x; *r* = 0.71, *p* = 0.022) and in genome coverage (22.5%; *r* = 0.86, *p* = 0.059). Porcine bocavirus (PBoV), a non-enveloped, single-stranded DNA virus of the *Parvoviridae* family which causes respiratory and gastrointestinal disease in young pigs and is associated with coinfections that exacerbate enteritis, was detected in 70% of samples, and showed significant enrichment in rPM (2.8x, *p* <0.001), and a strong increase in genome coverage (17.6%; *r* = 0.72, *p* = 0.006). Porcine sapelovirus (PSapeV), a non-enveloped, positive-sense single-stranded RNA virus of the *Picornaviridae* family, which causes gastroenteritis, pneumonia, and occasionally polioencephalomyelitis in pigs, was detected in 73% of samples, and showed slight positive enrichment in rPM (1.24x; *r* = 0.34), and a slight increase in genome coverage (3.6%; *r* = 0.14). Enterovirus G (EV-G), a positive-sense single-stranded RNA virus of the *Picornaviridae* family, which causes enteric disease and growth retardation in pigs, was detected in 65% of samples, and showed slight but consistent enrichment in rPM (1.23x; *r* = 0.62, *p* = 0.015), and a positive increase in genome coverage (7.7%; *r* = 0.32). Porcine kobuvirus (PKV), also known as Aichivirus C, is a non-enveloped, positive-sense single-stranded RNA virus in the family *Picornaviridae*, which is often detected in co-infections with other enteric viruses, was detected in 54% of samples, and showed strong enrichment in rPM (4.7x; *r* = 0.79, *p* = 0.023), and in genome coverage (9.2%; *r* = 0.76, *p* = 0.019). Porcine teschovirus (PTeV), a non-enveloped, positive-sense single-stranded RNA virus of the *Picornaviridae* family, which causes teschovirus encephalomyelitis characterized by paralysis, ataxia, and high mortality, was detected in 54% of samples, and showed strong enrichment in rPM (1.5x; *r* = 0.64, *p* = 0.032) and in genome coverage (9.9%; *r* = 0.76, *p* = 0.025). The most abundant targeted viral pathogen, Porcine astrovirus (PAstV), a non-enveloped, positive-sense single-stranded RNA virus of the family *Astroviridae*, was detected in 76% of samples and showed enrichment in rPM (1.12x; *r* = 0.25, *p* = 0.329) and in genome coverage (4.7%). Finally, we assessed enrichment of HPV18 oncogenes (E6/E7). MSSPE showed strong enrichment (2.9x; *r* = 0.91, *p* = 0.062) and a 4% increase in coverage of the E6/E7 loci. Since HeLa cells only have partial (∼67%), fragmented HPV-18 DNA integrated into their chromosomes, this relatively low increase is likely due to our primers exclusively targeting the HPV18 oncogenes (E6 and E7) rather than the whole genome.

Wilcoxon-ranked sum test assessed paired differences for fold change across samples. Adjusted p-value uses false discovery rate adjustment (Benjamini–Hochberg). **(b)** Genome coverage breadth increase per viral target. % coverage represents mean genome coverage breadth. Percent improvement in mean genome coverage values for each library prep method. Effect size is calculated as a rank-biserial correlation (r) from a paired Wilcoxon signed-rank test. **(c)** For each viral target, table shows the number of discordant detections where the virus was identified by MSSPE only or by mNGS only across matched sample–library preparation pairs. An odds ratio (MSSPE/mNGS) with 95% confidence intervals is reported.

Fifty-one swine slurry libraries were sequenced at an average depth of 31,357,075 reads (Interquartile Range (IQR): 4,187,471 – 51,273,014), with an average of 3,837,586 preprocessed reads (IQR: 2,761,058 – 4,711,943). Eight agricultural wastewater libraries were sequenced at an average depth of 44,012,813 reads (IQR: 20,635,710 – 76,384,922), with an average of 3,683,533 preprocessed reads (IQR: 2,899,278 – 4,113,938) (**Supplementary Table 1**). mNGS samples were sequenced at an average depth of 28,569,379 reads, with 3,384,868 preprocessed reads passing QC filters (IQR: 2,469,244 – 4,178,364), whereas MSSPE samples were sequenced at an average depth of 43,212,083 reads with 4,134,220 preprocessed reads passing quality control (QC) filters (IQR: 2,837,956 – 5,367,048) (**Fig. 3a**).

**Figure 3.**
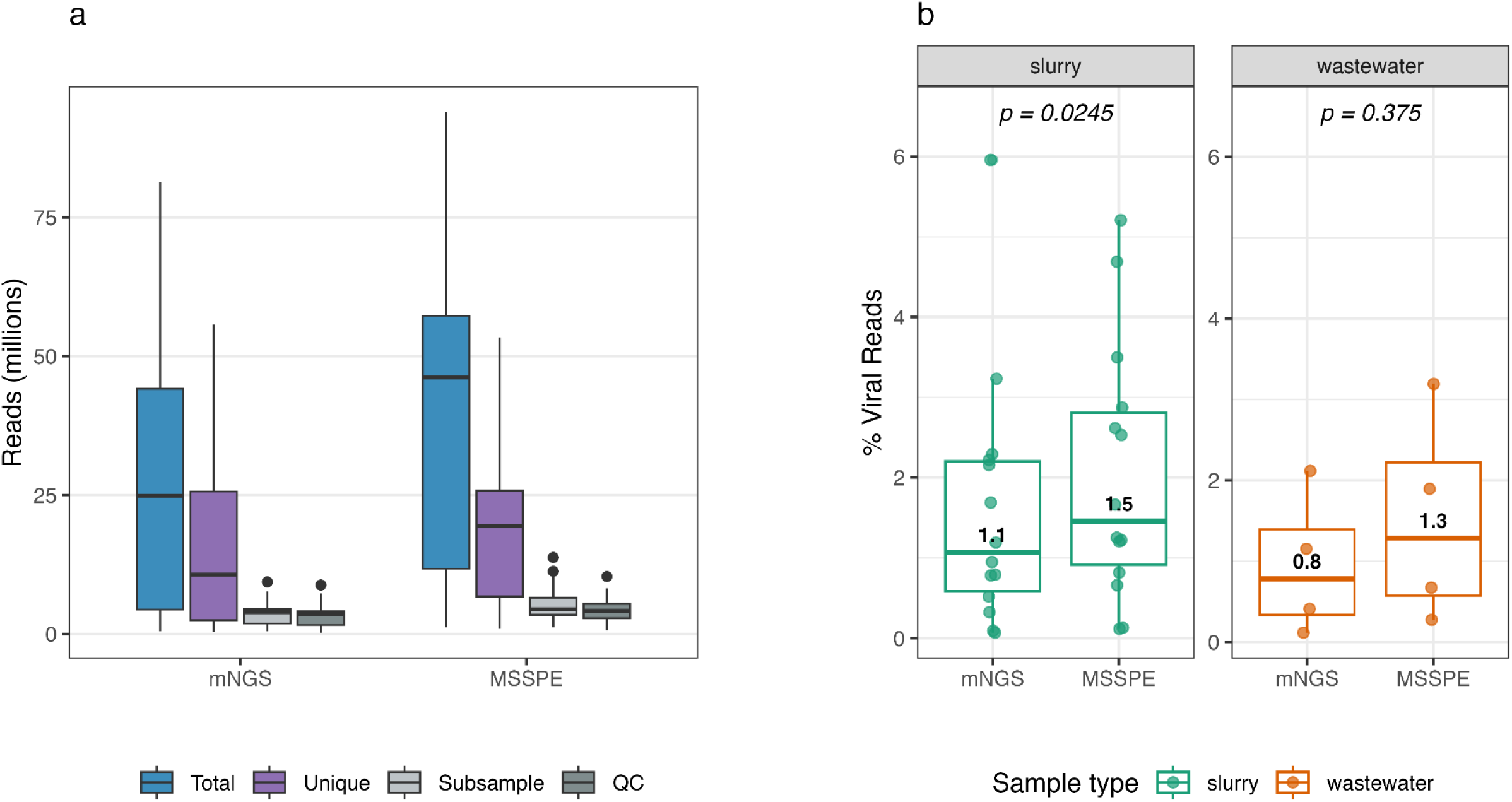
**(a)** Overall sequencing performance by library preparation method (MSSPE or untargeted mNGS) according to mean total reads, mean unique reads, mean subsampled reads, and mean reads passing all QC filters. **(b)** Percentage of reads aligned to viral taxa for each library preparation method. Wilcoxon p-value compares paired samples’ non-phage viral percentages. Each point represents a paired sample viral percentage value. Median values shown above the median line.

While sequencing output was higher for MSSPE libraries, the difference in reads passing CZ ID’s QC filters, which are the only reads taxonomically classified, was not substantial. MSSPE libraries had a substantially higher proportion of reads aligned to both phage and non-phage viral taxa (mNGS: median 1.1%, IQR 0.44-2.15; MSSPE: median 1.8%, IQR 0.93-2.82, *p* = 0.004) (**Fig. 3b**). Our results show that the difference in proportion of viral reads remained after grouping libraries by sample type, and that slurry samples had a significantly higher proportion of non-phage viral reads in MSSPE libraries relative to wastewater samples (**Fig. 3b**).

Consistent with prior spiked enrichment studies^1^, MSSPE showed incidental enrichment in untargeted viral genera which contain sequence homology to our spike primer sequences (**Supplementary Fig. 1**). While we did not detect presence of FMDV in any experiments, the results showed enrichment of two other porcine viruses not initially targeted, PKV (median 1.6x, *p* = 0.13) and EV-G (median 2.8x, *p* = 0.46), both of which are members of the same viral family as FMDV (*Picornaviridae*), though the incidental enrichment was not significant. Additionally, PAstV (*Mamastrovirae*) showed non-significant incidental enrichment (median 1.8x, *p* = 0.16).

### 2.3 MSSPE shows promise for low sequencing depth

Deep sequencing of complex samples (including wastewater, pig caeca, and cattle feces) demonstrates that near-complete taxonomic composition can be recovered at 500,000–1 million reads per sample and that relative proportions of taxa classifications remain constant regardless of sequencing depth^25–27^. To determine whether our observed differences in enrichment could be attributed to sequencing depth, taxon-level read counts were downsampled to varying depths using random subsampling at the shallow-depth thresholds (100,000–1.5 million reads per sample). We modeled the probability of detecting our targeted viral taxa using a binomial generalized linear mixed-effects model, with library preparation and sequencing depth as fixed effects, and each viral target and sample as random effects (**Table 3**).

**Table 3.** Fixed and random effects from the generalized linear mixed-effects model of virus detection probability. Fixed effects show the estimated influence of library preparation method, sequencing depth (scaled per 100,000 reads), and their interaction on the probability of detecting a virus. Estimates are presented on the log-odds scale, with corresponding standard errors, z-values, p-values, odds ratios (OR = exp(β)), and 95% confidence intervals for the OR.

| Effect | Estimate | SE | z | p | OR | CI<br>25% | CI<br>75% | Percent<br>change |
| --- | --- | --- | --- | --- | --- | --- | --- | --- |
| Library<br>prep | 2.124 | 1.327 | 1.600 | 0.11 | 8.361 | 0.620 | 112.716 | 736.073 |
| Sequencing<br>depth | 0.207 | 0.008 | 25.115 | 0.00 | 1.230 | 1.211 | 1.251 | 23.047 |
| Library<br>prep *<br>Se-<br>quenc-<br>ing<br>depth | -0.081 | 0.011 | -7.333 | 0.00 | 0.922 | 0.903 | 0.943 | -7.755 |

As expected, sequencing depth strongly increased detection probability, with each increase in 100,000 reads increasing the likelihood of detecting target viruses by 23% (OR 1.23, CI 1.21-1.25 , *p* <0.001). MSSPE showed a trend towards increased detection, but the effect was not significant with extremely wide confidence intervals (OR 7.7, CI 0.64-92.96, *p* = 0.11). However, when adding an interaction between library preparation and sequencing depth, MSSPE did increase the probability of detecting target taxa at shallow depths. For each increase in 100,000 reads, the odds of detecting a targeted virus in MSSPE libraries increases 6-10% *less* than mNGS (OR 0.92, CI 0.90–0.94, *p* <0.001) (**Fig. 4c-e**). This indicates that MSSPE is more efficient at lower sequencing depths, and has higher odds of detecting low-abundance viruses earlier than mNGS, requiring fewer reads to identify targeted viral taxa. We observed substantial virus- and sample-specific variability, with variance of σ² = 22.16 (SD = 4.7) for viral taxon and σ² = 8.00 (SD = 2.8) for sample, indicating that enrichment varies substantially by viral species and, to a lesser extent, across samples. The model’s explanatory power appears to be dominated by random effects, consistent with the substantial observed heterogeneity in enrichment and genome coverage across viral targets and samples.

**Figure 4.**
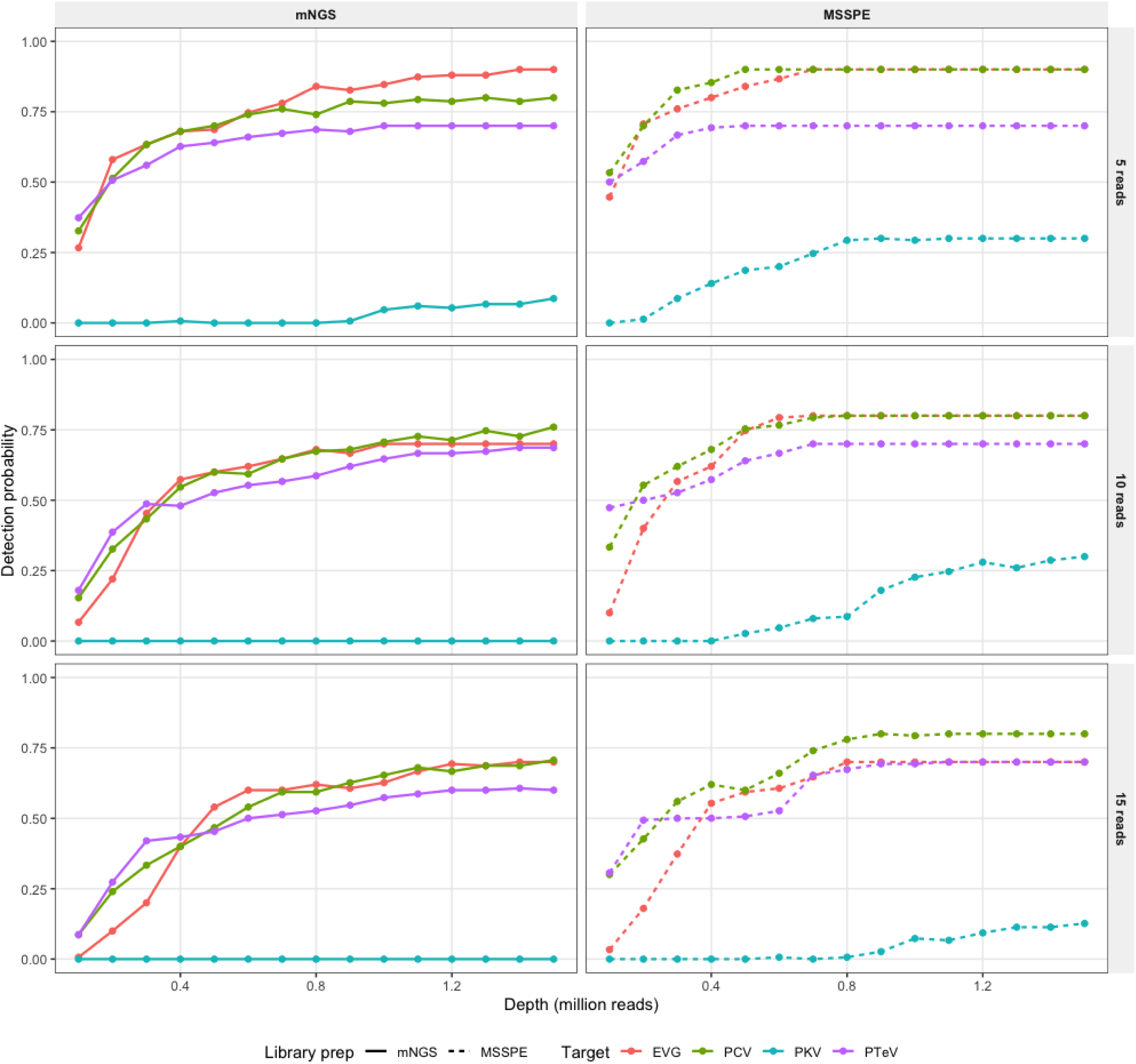
Proportion of rarefaction replicates where viral taxa met or exceeded defined detection thresholds (reads >= 5, 10, 15). Detection probability was then summarized as a function of sequencing depth and library preparation. Viral target species identified by line color. MSSPE replicates identified by dashed line. mNGS replicates identified by solid line.

Subset of targets shown for clarity. All viral targets shown in Supplementary Figure 2.

### 2.4 MSSPE and mNGS captured similar microbial diversity

To evaluate whether spiked primer enrichment biased microbial recovery, we compared sample composition, species diversity, and viral composition between mNGS and MSSPE. Across both methods, a total of 93 non-phage viral genera were detected, consisting of 502 unique species of non-phage viruses. Our results show that 6% of non-phage viral species were known to infect pigs, chickens, or humans. Consistent with previous metagenomic studies of wastewater^19,28,29^, the vast majority of reads aligned to bacteria (72.6%), followed by eukaryotes (17.0%), viruses (5.6%), and archaea (4.9%) (**Fig. 5a**). The distribution of biological groups within samples was largely similar between mNGS and MSSPE, with MSSPE displaying slightly lower proportions of archaea (mNGS: 6.7%, MSSPE: 3.4%) and slightly higher proportions of eukaryotes (mNGS: 15.3%, MSSPE: 18.3%), while proportion of viral reads was nearly identical (mNGS: 5.5%, MSSPE: 5.7%). However, sample composition differed substantially by source, with slurry samples retaining a significantly higher proportion of viral taxa (slurry: 6.5%, wastewater: 1.7%), bacterial taxa (slurry: 76.7%, wastewater: 53.9%), and a significantly lower proportion of eukaryotic taxa (slurry: 11.6%, wastewater: 41.5%) (**Fig 5a,5b**).

**Figure 5.**
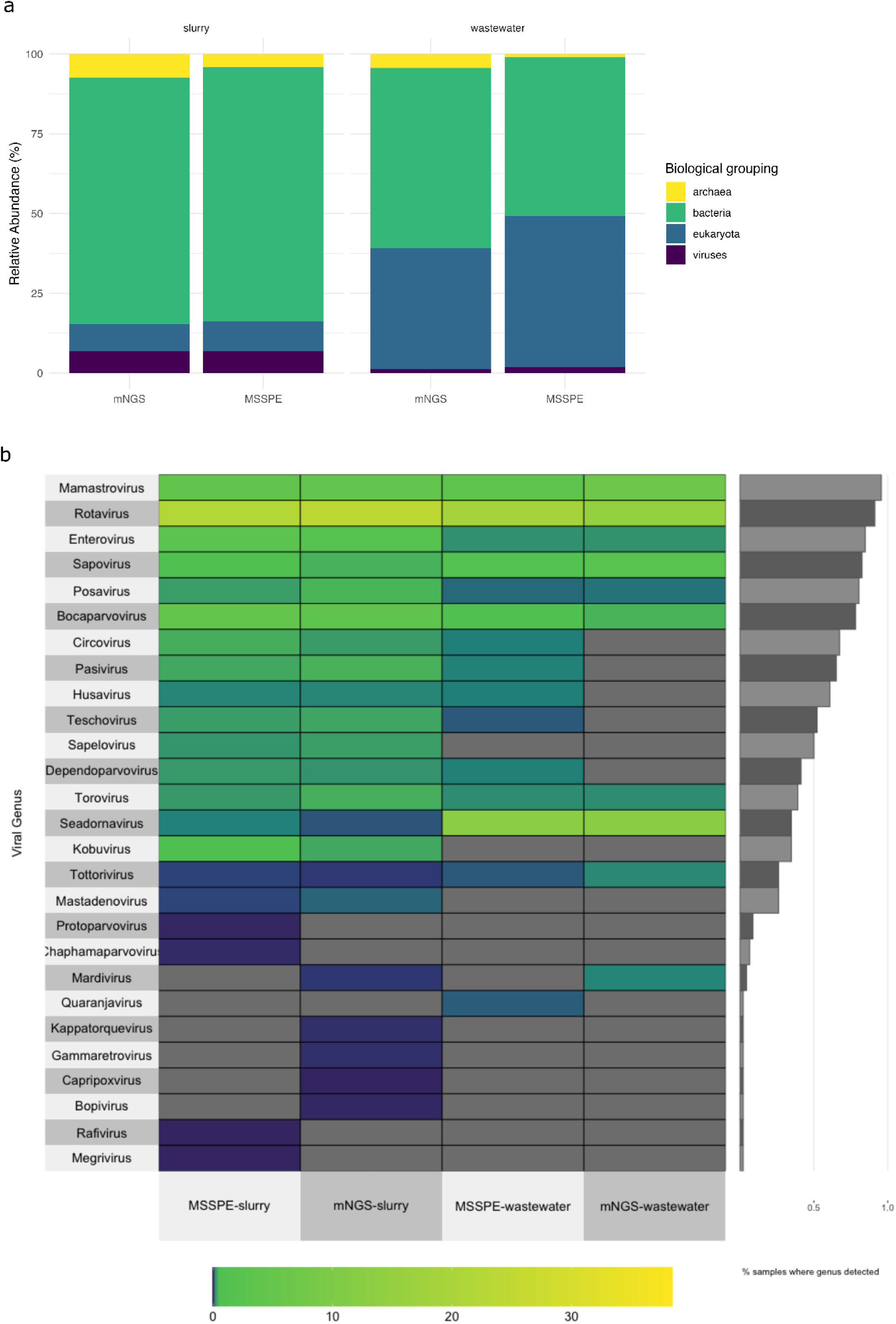
**(a)** Sample composition via relative abundance (%) per library prep method and sample type **(b)** Viral composition heatmap comparing relative abundance (%) of vertebrae-infecting viral genera between MSSPE and mNGS. Lighter heatmap colors indicate higher relative abundance. Grey color on the heatmap indicates taxa that were not detected for a given library prep or sample type. Barchart on the right y-axis shows the proportion of samples where genera had at least one species detected.

**Figure 6.**
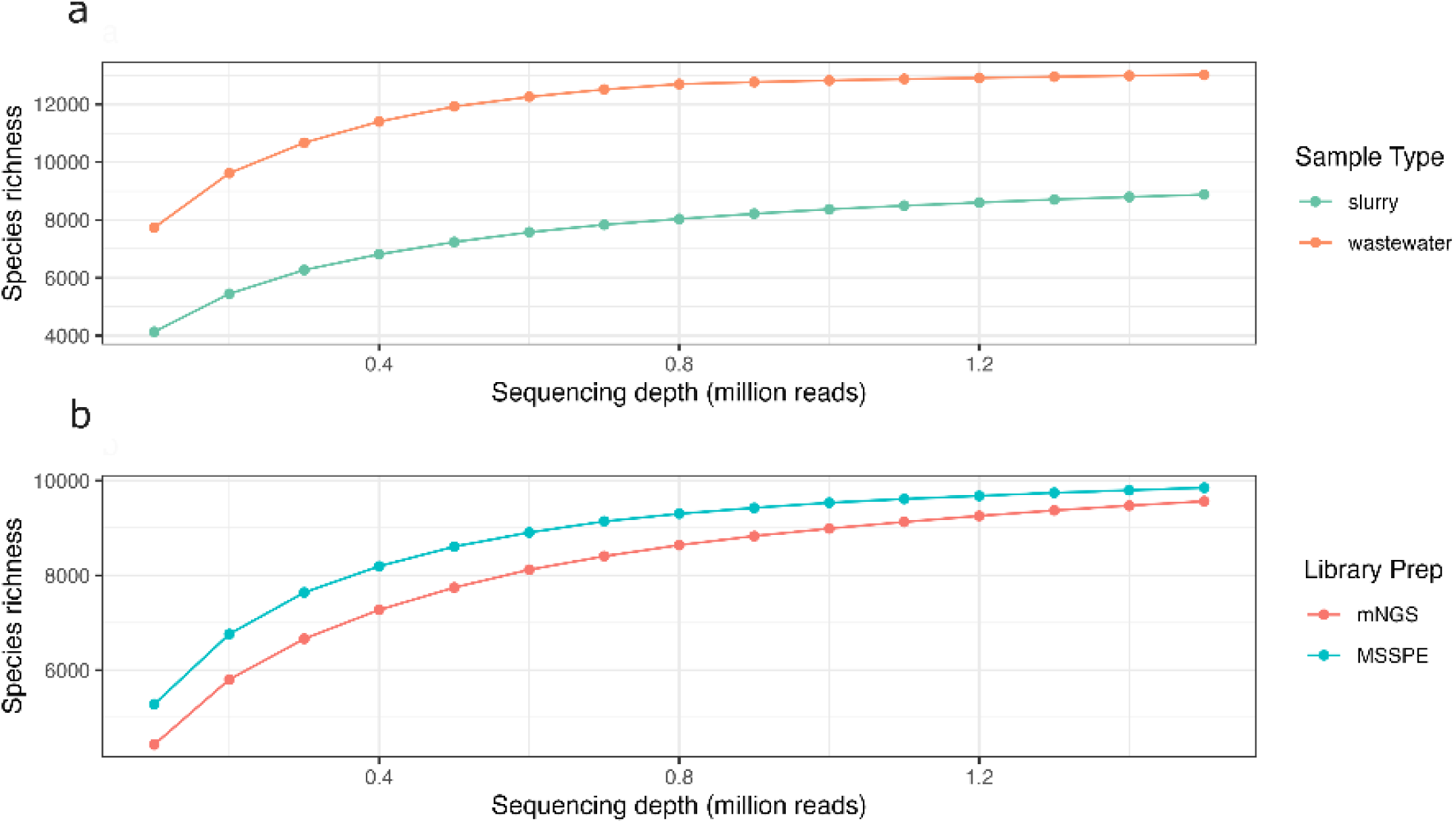
**(a)** Species richness at increasing sequencing depths per sample type. Slurry represents a mixture of pig feces, urine, and water used to flush manure from pig housing. Wastewater represents downstream agricultural effluents from slurry runoff housed in a pond. **(b)** Species richness at increasing sequencing depths per library preparation method.

**Figure 6.**
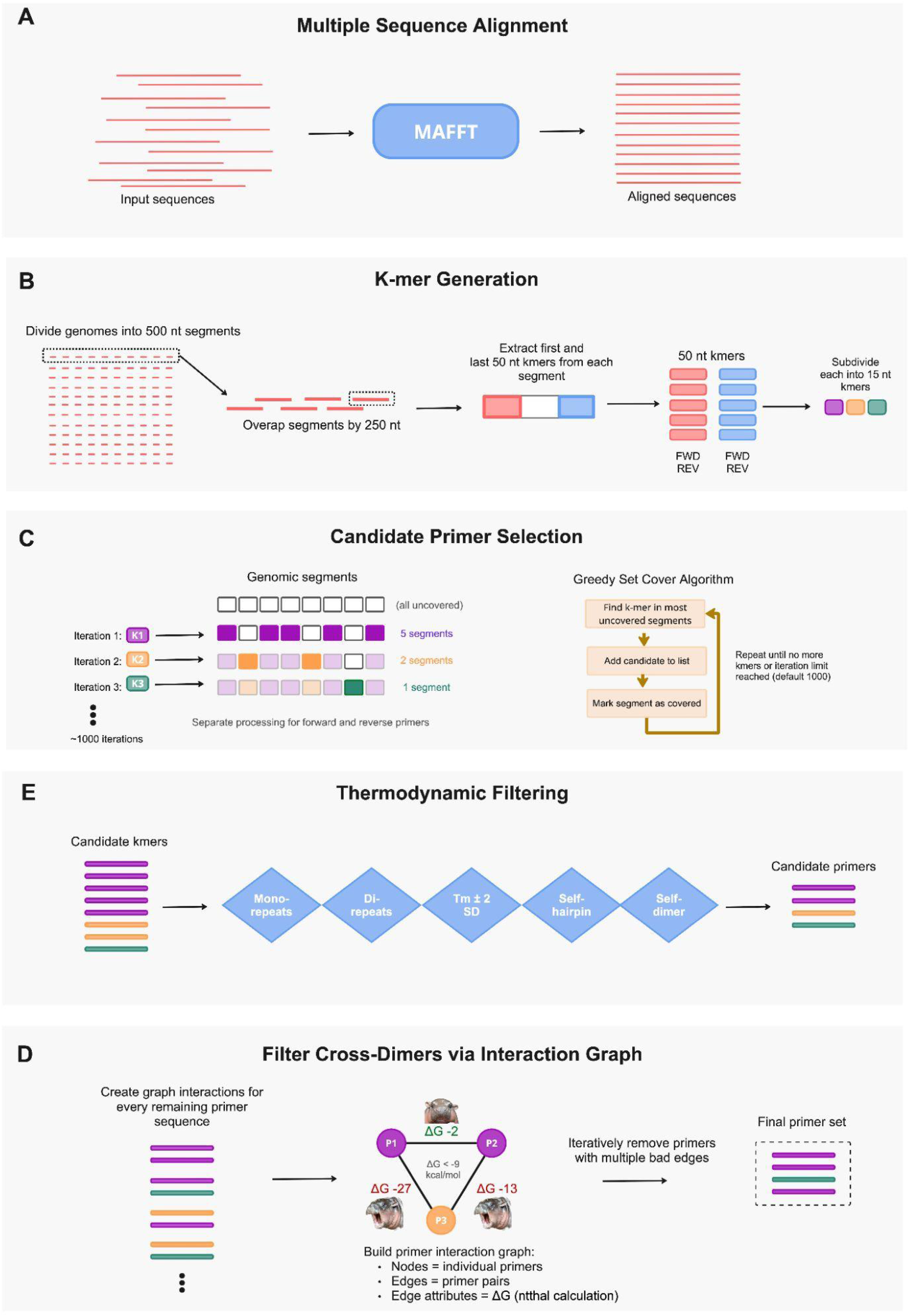
Primer design algorithm used to automatically generate spiked primers. Algorithm is executed for each viral target separately. **(a)** Full-length genomes are multiple-sequence aligned using MAFFT. **(b)** The aligned genomes are then partitioned into overlapping segments (default 500 nt windows with 250 nt overlap). **(c)** For each segment, k-mers (default 15 nt) are generated from both the forward and the reverse-complement k-mers from terminal ends of segments. K-mers are ranked by the number of distinct reference sequences in which they occur. **(d)** Candidates are filtered by sequence composition (removal of homopolymers or repeats) and thermodynamic properties. We compute melting temperature (T_m_) and Gibbs free energy changes (ΔG) for secondary structure formation (e.g. hairpins, self-dimers). **(e)** All-vs-all pairwise ΔG values were calculated using ntthal, and interactions were represented as a graph where nodes correspond to individual primers and edges store pairwise ΔG values. Primers involved in multiple strong cross-dimer interactions (ΔG < -9 kcal/mol) were iteratively removed.

Sample type primarily drove differences in microbial community diversity between MSSPE and mNGS across three metrics. Diversity of wastewater samples was not significantly affected by library preparation method, whereas slurry samples showed significantly higher species richness when prepared with MSSPE (slurry: *p* = 0.02, wastewater: *p* = 0.62) (**Supplementary Table 2, Supplementary Figure 3**). The samples with the most diverse microbial community were collected from agricultural wastewater retention ponds on the farms (**Supplementary Table 2**), while the samples with the highest proportion of viral reads were collected from slurry pits (**diversity: Supplementary Table 2, viral reads: Fig. 3b**). While MSSPE and mNGS had similar numbers of distinct viral genera in both slurry and wastewater samples, several low abundance genera were detected exclusively by one method or the other (**Fig. 5b**). Specifically, *Protoparvovirus, Chaphamaparvovirus, Rafivirus*, and *Megrivirus* were only detected in MSSPE libraries from slurry samples, while *Bopivirus, Mardivirus, Kappatorquevirus*, and *Gammaretrovirus* were only detected in mNGS libraries from slurry samples.

Rotavirus was the most abundant genus of RNA virus in the samples, followed by *Orthopicobirnavirus, Kobuvirus, Enterovirus, Seadornavirus, Sapovirus, and Pasivirus* (**Fig. 5b**, **Supplementary Table 3**). *Bocaparvovirus* was the most abundant genus of DNA virus, followed by *Circovirus*. Out of 59 libraries sequenced, *Rotaviruses* were found in 91% of libraries, *Enteroviruses* 85%*, Sapoviruses* 83%*, Bocaparvoviruses* 78%*, Circoviruses* 67%*, and Kobuviruses* 35% (**Fig. 5b, Supplementary Table 3**). Relative to mNGS, MSSPE libraries showed a 4-fold increased odds in detecting *Seadornaviruses,* though confidence intervals were wide (0.85-18.84), a 1.8-fold increased odds for *Kobuviruses* (0.60-5.37), a 1.3-fold increased odds for *Bocaparvoviruses* (0.94-1.88), a 1.6-fold increased odds for *Circoviruses* (0.80-3.07), and a 1.6-fold increased odds for *Rotaviruses* (1.08-2.38). Overall, the dominant viral taxa detected were enteric RNA viruses of the *Picornaviridae* and *Astroviridae* families, reflecting the high prevalence of fecal–oral transmission and environmental persistence in intensive pig farming.

To determine whether differences in diversity could be attributed to sequencing depth, taxon-level read counts were downsampled to varying depths using subsampling at the shallow-depth thresholds (100,000–1.5M reads per sample). We modeled species richness using a negative binomial generalized linear mixed-effects model, with library preparation, sequencing depth, and sample type as fixed effects, and each viral target and sample as random effects (**Supplementary Table 5**). Species richness was strongly influenced by sample type, with slurry samples exhibiting ∼46% lower richness than wastewater samples after accounting for depth and library preparation (OR 0.544, 95% CI 0.390–0.759, *p* < 0.001) (**Fig. 6a)**. Sequencing depth had a small positive effect on richness (∼3.2% increase per 100,000 reads), with a stronger depth–richness relationship in slurry samples (OR 1.015, 95% CI 1.011–1.018, *p* < 0.001). In contrast, MSSPE modestly increased richness (∼20%; OR 1.199, 95% CI 1.152–1.248, *p* < 0.001). However, MSSPE did not meaningfully modify the effect of sequencing depth (OR 0.988, 95% CI 0.984–0.993, *p* < 0.001) or sample type (OR 0.962, 95% CI 0.920–1.006, *p* = 0.090), suggesting that at lower sequencing depths, MSSPE does not substantially bias sample composition, even in slurry samples (**Fig. 6b**).

## 3. DISCUSSION

This study presents the development and preliminary validation of an enriched metagenomic assay, optimized for wastewater collected from rural livestock farms in Southeast Asia. MSSPE retained the sensitivity of untargeted metagenomics, as measured by species diversity, and increased the ability to detect low-abundance viral targets, as measured by viral enrichment, without significantly increasing sequencing depth. Our approach demonstrates that spiked primer enriched metagenomics can be successfully adapted for complex environmental samples in resource-constrained settings to detect high-risk priority DNA and RNA viral pathogens, including known zoonoses with important implications for public health.

A key contribution of this work is the development of the open-source primer design algorithm, *Open MSSPE Design*, optimized specifically for efficient creation of spiked primers for MSSPE protocols. Using this tool, we produced 15 primer sets targeting diverse DNA and RNA viral pathogens, with *in silico* validation confirming robust coverage across all targets, and laboratory experiments validating enrichment performance for at least nine targets in real-world samples. The pipeline’s ability to automatically assess secondary structure stability across pooled primer sets provides a valuable tool that may extend beyond MSSPE and wastewater workflows, and could be adapted for researchers designing primers for targeted PCR, amplicon sequencing, or other primer-based applications.

One of the primary benefits of metagenomics for outbreak prevention is the capability to detect all nucleic acids in a sample, including priority, emerging, and novel pathogens. However, sequencing low abundance viral pathogens from high background complex sample types, such as wastewater, at adequate depth to robustly detect priority pathogens has previously limited the application of mNGS outside of well-funded research settings. Our enrichment approach addresses this fundamental limitation by increasing the likelihood of identifying priority pathogens, while maintaining the broad detection capabilities that make metagenomics valuable for outbreak surveillance. Notably, our MSSPE-based assay added no additional time to metagenomic library preparation when compared to mNGS, while requiring only the purchase of synthesized primers beyond standard mNGS reagents. Considering the added time and cost for viral capture library preparation protocols^21^, MSSPE may be an affordable alternative for LMICs to enrich wastewater samples for metagenomics. However, considering substantially higher enrichment reported by hybrid capture methods (>100x)^16^, MSSPE could be considered a supplementary or preliminary screening approach, helping inform the decision to procure extremely expensive hybrid capture probes (estimated cost of commercial kits in Thailand ∼US$18,500 for 12 reactions). MSSPE could also be combined with hybrid capture or multiplex PCR to further increase genome coverage, as prior studies have achieved 25-80% increases in clinical samples^1^.

Additionally, the upper limit for the number of taxa that can be simultaneously enriched via MSSPE has yet to be experimentally determined. In contrast, commercially available hybrid capture products report enrichment of over 3,000 viral taxa known to infect vertebrates (though some evidence suggests that broader panels which target more viruses might actually have lower sensitivity in detecting lower abundance viruses^30,31^). The incidental enrichment we observed may be related to highly conserved regions across species in the same family as the spiked primers. This enrichment could be due to the genome-wide design of our spiked primers, resulting in amplification of highly-conserved regions, or may reflect the diversity of reference genomes used in primer design. While the off-target amplification was unintentional, these findings suggest the potential for MSSPE to broadly target higher order taxon levels, rather than specific species taxa, which may be a utility for enriching entire viral families or genera^24^.

Our results demonstrate significant enrichment across most viral targets, though the magnitude of enrichment was lower than reported in previous MSSPE studies, which were conducted with low-background clinical samples. This reduced enrichment efficiency likely reflects fundamental differences between clinical and environmental sample matrices, as well as several interrelated factors that warrant consideration for future investigation.

We identified several factors likely to contribute to the heterogeneous enrichment performance we observed across viral targets. Genome structure may play a role, as segmented viruses like influenza present distinct challenges for primer design compared to non-segmented genomes, potentially requiring higher primer densities to ensure adequate coverage across all segments. Primer density itself—measured as primers per kilobase of genome—varied across our targets (**Table 1**) and may directly influence enrichment efficiency, with optimal densities likely differing between short genomes and large genomes. Viral abundance in samples constrains detection probability, as evidenced by our depth-dependent detection models showing that even enriched libraries require minimum target concentrations. RNA integrity and fragmentation in wastewater likely reduce primer binding opportunities, particularly for targets with degraded genomes. Differences in capsid and envelope resilience may explain why some viruses persist better in harsh environmental conditions, affecting detection regardless of enrichment method. The complex microbial communities present in swine slurry or agricultural wastewater create competition for primer binding sites and reverse transcription resources. It is possible that long-read sequencing technology could overcome some of these challenges^32^, and should be explored in future research. However, it is also possible that our primer design algorithm, while validated *in silico* via simulated coverage, may not be optimally configured for the specific conditions present in wastewater samples. Indeed, while our primer design algorithm can design primers of any length, we were limited by supplier restrictions to a minimum primer length of 15 nucleotides. Prior MSSPE studies used the same supplier to synthesize 13 nucleotide primers, and it is possible that the shorter primers are more optimal for MSSPE-based amplification.

For rapidly evolving viruses, primer design presents specific challenges that must be balanced against the method’s advantages. MSSPE requires prior knowledge of target sequences, which could be viewed as limiting; however, designing primers against highly conserved genomic regions may suffice to detect even novel variants within known viral families. This represents a middle ground between purely untargeted metagenomics and highly specific amplicon approaches. The method’s reliance on sequence conservation means it may miss entirely novel viral families with no sequence homology to known pathogens, a limitation inherent to any targeted approach. Additionally, enrichment magnitude remains substantially lower than hybrid capture methods (2-fold vs. >100-fold), positioning MSSPE as optimal for monitoring known or emerging threats rather than pathogen discovery. To maintain effectiveness as viruses evolve, primer panels should be updated periodically by informed phylogenetic surveillance data and regular in silico validation against contemporary sequences. We suggest establishing review cycles—ideally annually or following major outbreak events—during which primer sets are evaluated against updated sequence databases and redesigned as needed to maintain coverage of circulating strains, a process greatly facilitated by the automated, open-source nature of our primer design pipeline.

Substantial increases in rPM did not always correlate with increases in genome coverage. Several targeted viruses (PKV, PCV, PBoV, PTeV) showed strong increases both in rPM and genome coverage, whereas EV-G showed non-significant increases, despite strong enrichment. Finally, while we did not enrich for any enveloped viruses, we did detect several untargeted enveloped viral taxa in both MSSPE and mNGS samples, including the swine pathogen *Torovirus suis* (Porcine torovirus) and the emerging Orthomyxovirus genus *Quaranjavirus*^33–36^. Future research should investigate why particular viruses seem to respond more strongly to MSSPE than others, and whether enveloped viruses can be enriched via MSSPE in wastewater.

Our results suggest that MSSPE libraries may retain a higher proportion of viral reads. Similar to MSSPE of clinical samples, we hypothesize that primer spiking can increase the relative proportion of viral fragments in the library, even in the presence of random hexamers, which still allow for priming of untargeted templates. We also found that slurry samples, which contained more solids than our agricultural wastewater samples, had a significantly higher proportion of viral taxa. This is largely consistent with recent evidence from human wastewater studies suggesting that viruses, in particular enveloped viruses, preferentially partition to settled solids rather than influent wastewater^37–39^. Interestingly, slurry samples showed significantly higher species richness when prepared with MSSPE. While it is possible that MSSPE biases sample types dominated by solid material, we did not quantify the differences in organic matter between sample types, and the precise mechanism explaining why slurry samples showed disproportionate influence by spiked primer enrichment remains unclear.

MSSPE may be more efficient for detecting low-abundance viruses with fewer reads. Our experiment showed that MSSPE boosted viral detection at lower sequencing depths, while mNGS libraries required a significant increase in sequencing depth in order to achieve the same probability of detection. The magnitude of enrichment per viral taxa was highly heterogeneous, underscored by our model’s high variance largely due to viral species and between-sample differences. Sequencing depth increased detection odds by ∼23% per 100,000 reads across both methods, though MSSPE showed 8% less increase per unit depth than mNGS (OR 0.92), indicating higher baseline efficiency at shallow depths.

Beyond sequencing performance, our methodology demonstrates the practical feasibility of implementing metagenomic surveillance in resource-limited agricultural settings. The use of simple grab sampling techniques with readily available collection tools successfully captured a diverse microbial community, consistent with previous studies of grab sampling^40,41^, while requiring significantly fewer resources than composite sampling. However, grab samples may not fully capture the spatial and temporal variability in highly heterogeneous environments. Future research should explore comparisons between grab sampling and other more alternative sampling methods. The integration of PEG precipitation for sample concentration with MSSPE represents a practical approach for processing and enriching complex environmental samples.

The combination of open-source computational tools, simplified sampling protocols, and cost-effective laboratory methods creates a scalable approach that can be adapted to diverse geographic and economic contexts. Our research context is a resource-limited veterinary lab in northern Thailand with lab staff who, prior to this study, had never prepared libraries for metagenomics, nor worked with RNA or performed cDNA synthesis. Nevertheless, the requirements for biosafety, reliable cold chain logistics, consistent electricity supply, and skilled personnel may limit implementation in some extremely resource-constrained settings where surveillance is most needed.

## 4. CONCLUDING REMARKS

This work represents a pilot application of spiked primer enriched metagenomics for surveillance of high-risk pathogens in complex sample types. The successful adaptation of MSSPE to these challenging sample types demonstrates the flexibility of the approach and its potential for expansion to other surveillance applications. The development of a surveillance approach specifically designed for resource-limited settings addresses a critical gap in global pathogen surveillance infrastructure. To enable routine deployment of MSSPE in LMICs, future work should rigorously define limits of detection, expand panels to additional taxa including bacteria and eukaryotic parasites, evaluate performance across diverse clinical and environmental sample types, and benchmark turnaround times and per-sample costs against regional timeliness^42^ and infrastructure constraints to further reduce the overall cost of metagenomic surveillance.

## 5. MATERIALS AND METHODS

### 5.1 Overview of MSSPE assay development

From September 2024––July 2025, 21 swine slurry samples and 4 agricultural wastewater samples from five mixed-livestock farms in northern Thailand were collected. For the purposes of this study, *swine slurry* refers to a mixture of pig feces, urine, and water used to flush manure from pig housing, typically collected from pit, well, or gutter systems in swine production facilities^43^. In contrast, *wastewater* encompasses a broader category of liquid waste, including agricultural effluents that are downstream from the slurry runoff often housed in a pond.

We sought field sample collection protocols that balanced molecular efficiency and affordability, based on our hypothesis that protocols^44^ from human wastewater surveillance systems can be adapted to swine slurry in rural settings, building on the work by Ramesh and Bailey et al. (2021)^45^. Aligning with our participatory research goals, we also sought agreement with local farm owners prior to visiting their farms so as to not significantly disturb their farm operations, and received approval on the condition that our methods were entirely non-invasive. We collected two samples using decontaminated plastic cooking ladles from the same source from five locations around each farm including slurry gutters, slurry pits, mixing wells, and effluent ponds. We selected a simplified polyethylene glycol (PEG) precipitation method^46–49^, over expensive single-use ultrafiltration membranes, based on work of Barnes et al. (2023) for its affordability and relatively high yield as demonstrated experimentally with grab sampling in an LMIC setting.

After concentration and nucleic acid extraction, we conducted paired experiments in which two laboratory staff worked in parallel on each sample to compare enrichment performance between MSSPE and mNGS. Initially, we pooled both reverse and forward primers together into a single reaction, for eight (8) targeted DNA and RNA pathogens: *Asfivirus haemorrhagiae* (African swine fever virus)*, Pestivirus suis (*Classical swine fever virus)*, Aphthovirus vesiculae* (Foot-and-mouth disease virus), *Henipavirus nipahense* (Nipah virus), *Alphainfluenzavirus influenzae* (Influenza A virus), Porcine circovirus, *Betaarterivirus europensis* (Porcine reproductive and respiratory syndrome virus), and a process control for partially integrated *Alphapapillomavirus 7* (Human papillomavirus 18). These samples were sequenced on an Illumina Nextseq 2000 with an average sequencing depth of 7.7 M reads per sample.

Next, in part because our research team had limited access to target RNA for true positive controls, we sought to validate our assay against a greater number of targets which were likely to be present based on the initial experiment. We performed a second experiment on the same samples (both swine slurry and agricultural wastewater) but with seven new spiked primer sets targeting seven (7) new viral pathogens: *Aichivirus A* (Porcine kobuvirus*), Enterovirus geswini* (Enterovirus G), *Sapelovirus anglia* (Porcine sapelovirus), *Sapovirus sapporoense* (Porcine sapovirus), *Mamastrovirus suis* (Porcine astrovirus), *Teschovirus asilesi* (Porcine teschovirus) and *Bocaparvovirus ungulate* (Porcine bocavirus), in addition to a process control for the E6 and E7 HPV18 genes that are transcribed in HeLa cells. Finally, we collected a new set of samples from different farms and prepared libraries using all spiked primer sets targeting all 15 viruses. For greater specificity (as well as cost-savings in primers) the number of reference genomes in designing spiked primers included only references which shared at least 75% similarity to each other. These samples were sequenced at a significantly greater depth (∼53 M reads per sample) with an Illumina Novaseq X.

Downstream bioinformatic analysis focused on viral pathogens. Viral taxa are usually the least abundant phylum found in wastewater^16^, making them the most likely candidate to benefit from enrichment. Sequence contigs were generated using reads from each sample and classified using the taxonomic classification cloud-based software Chan Zuckerberg ID (CZ ID). Contigs were aligned to reference genomes for species confirmation using NCBI BLAST and Geneious Prime (v2025.1.3). Viruses pathogenic to humans, pigs, and/or chickens were flagged using the open source host-pathogen reference database Virus-Host DB^50^.

### 5.2 Site enrollment

Sample sites were selected by engaging local villages in Chiang Mai and Lamphun provinces in Thailand. The location of these villages border protected forest land, where abundant wildlife frequently venture outside of their habitat and come into contact with the local community and livestock. These farms are existing participants in our participatory disease surveillance projects. These local community members are already effective disease detectives, receptive to skills training with high levels of digital literacy, and are accustomed to outside researchers visiting their farms. We built upon our existing positive relationship with these local communities to maximize participation and impact of our capacity building activities.

### 5.3 Sample collection

A total of 25 samples were collected from 5 sites in Lamphun and Chiang Mai provinces. Two duplicate 30–40 mL samples were collected from each collection location using a decontaminated single-use plastic cooking ladle. Ladle decontamination protocol was adapted from the South African Medical Research Council’s Wastewater Sampling Guide^51^.

Briefly, we scrubbed ladles with a wire brush to remove large particles. Then we rinsed ladles with deionized or distilled water. Ladles were then soaked in a 10% bleach solution for 5 minutes. Ladles were washed a final time with deionized or distilled water, and then covered in plastic wrap after drying until use. The sampler used a new ladle for each collection along with full PPE (disposable full-body coveralls, N95 mask, face shield, non-powdered gloves). A digital record using the mobile data collection tool KoboToolbox (kobotoolbox.org) complemented each sample. Sample metadata included photos of the collection location and sample barcode, time/date of collection, name of sampling site, latitude/longitude coordinates, sample volume, and number and type of animals at site.

### 5.4 Sample processing

We adopted polyethylene glycol (PEG) precipitation methods based on work by Barnes et al^48^. Briefly, we added 3 g of PEG 8000 (Sigma Aldrich P2139-500G) and 0.68 g of NaCl (RCI Labscan AR1166-P1KG) to a 30 mL grab sample in a 50 mL Falcon tube (Appendix 1). We manually shake by hand for 30 seconds or until the PEG and NaCl is dissolved. Up to 20 samples are then spun at 1200 g, 4 °C (1200 g; range from 800–2000 g) for 2 hours. After centrifugation we discard supernatant trying to remove as much liquid as possible without dislodging the pellet and then add 200 uL sterile 1x PBS (pH 7.4, Merck P4417-100TAB) to the pellet and vortex for 2 minutes. Samples were then transferred to (locking) 1.5 mL Eppendorf tubes. A 2X DNA/RNA Shield stabilization solution (Zymo Research) is added to samples at 1:1 ratio, and then mixed vigorously by vortexing. Samples were mixed with DNA/RNA Shield within 4 hours of sample collection to minimize viral RNA degradation. DNA/RNA Shield was added after sample concentration, rather than using 1:1 ratio with original sample volume, to reduce costs. Samples are stored at −20 °C prior to DNA and RNA extractions.

### 5.5 DNA/RNA extraction

For samples which contained an excess of solids, making it difficult to pipet, we homogenized the samples with Zymo Research Bashing Beads (Zymo Cat. #S6012-50) by vortexing for 1 minute followed by 1 minute incubation on ice for a total of 3 cycles using a high speed benchtop homogenizer. Then, we spun the sample to pellet the cell debris and bashing beads, and transfer the supernatant to a new tube. After incubation, we vortex the samples and centrifuge at max speed for 2 minutes to further pellet debris, and then transfer the cleared supernatant to a new nuclease-free tube.

For all samples, we performed a proteinase k treatment to optimize viral purification. Proteinase k treatment digests proteins to eliminate contamination from nucleic acid preparations, and inactivates nucleases which might degrade RNA. Briefly, for 400 uL of reagent/sample mixture, we added 10 uL Proteinase K and mixed thoroughly. Then we incubated at room temperature (20-30 °C) for 30 minutes. Then we added a 1:1 lysis buffer before purifying with the Zymo Pathogen Magbead kit.

We performed parallel DNA and RNA extractions with DNAseI digestion on column on all samples and nuclease-free water negative controls (Sigma Aldrich W4502-1L) using the Zymo Quick DNA/RNA Microprep Plus kit according to manufacturer instructions. After extraction, we performed quality control (QC) measures to assess the amount and integrity of input nucleic acid prior to library preparation using a Qubit Fluorometer 4 and a Nanodrop. RNA extractions were then stored at -80 °C prior to library preparation. DNA extractions were stored at -20 °C due to space constraints.

### 5.6 Spiked primer enrichment

#### 5.6.1 Open-source primer design algorithm

To design virus-specific spiked primers, we developed *Open MSSPE Design*, an automated and configurable Rust-based pipeline for designing virus-specific spiked primers, building on the MSSPE method described by Deng et al (2020) and introduces improvements in automation, candidate selection, and thermodynamic modeling **(Fig. 6)**. The pipeline performs the following steps:

1. <u>Sequence alignment and segmentation</u>. First, full-length viral genomes were retrieved from NCBI GenBank and multiple-sequence aligned using MAFFT v7.388 with default settings (algorithm = “Auto”, scoring matrix = 200 PAM/k = 2, gap open = 1.53, offset = 0.123), to identify conserved regions across strains, as in the original method. The aligned genomes are then partitioned into overlapping segments (default 500 nt window size; 250 nt overlap).
2. <u>K-mer Generation.</u> From each segment, terminal regions (default 50 nt from each end) are extracted, and all possible 15 nt k-mers are enumerated from these regions. Both forward k-mers and reverse-complement k-mers from the opposite segment end are collected as distinct candidates, allowing conserved target sites to be identified regardless of orientation within the alignment, consistent with bidirectional primer strategies used in MSSPE. Each candidate k-mer was indexed in a hash map tracking its occurrence across genomic segments, with conservation approximated by the number of segments containing each k-mer.
3. <u>Primer selection process.</u> Primer candidates were selected using a greedy set cover algorithm in which, at each iteration, the k-mer appearing in the most uncovered segments was selected, and those segments were marked as covered and excluded from subsequent iterations. This process was repeated for a user-defined number of iterations (default: 1000) or until no additional high-frequency k-mers remained, prioritizing broad genome coverage while limiting redundant targeting of overlapping regions.
4. <u>Thermodynamic characterization and filtering.</u> Following candidate selection, primers are characterized thermodynamically and then filtered by sequence composition and thermodynamic properties. We compute melting temperature (T_m_) and Gibbs free energy changes (ΔG) for secondary structure formation (hairpins, self-dimers, cross-dimers) using Primer3’s nearest-neighbor model. Primers with T_m_ outside a user-specified tolerance (default ± 2 SD from the mean) or with ΔG values more negative than a threshold (default -9 kcal/mol) are excluded.
5. <u>Cross-dimer compatibility graph analysis.</u> To ensure primers in the final panel would not form strong cross-dimers, all-vs-all pairwise ΔG values were calculated using ntthal. These interactions were represented as a graph where nodes correspond to individual primers and edges store pairwise ΔG values. Primers involved in multiple strong cross-dimer interactions (ΔG < -9 kcal/mol) were iteratively removed, prioritizing primers with the highest number of problematic interactions, until all remaining primers showed acceptable compatibility. All ΔG interactions are evaluated across the full primer set to ensure compatibility.

The final output comprises a set of primers that provide broad coverage of the target viral genome alignment while satisfying thermodynamic stability and compatibility constraints. Key parameters are configurable via a command-line interface, including the number of iterations, T_m_ range, ΔG cutoff, and segment and window size, allowing tuning for different viruses or design priorities. The source code is openly available at: https://github.com/opendream/open-msspe-design/

#### 5.6.1 Viral primers optimized for MSSPE

We used *Open MSSPE Design* (v1.0) with default parameters except for: kmer-size=15, min-tm=30, max-tm=60. Spiked primer sequences were validated *in silico* before synthesis using Geneious Prime (v2025.1.3) for optimal binding efficiency and amplification of target consensus genomes. Spiked primers were synthesized by Integrated DNA Technologies (IDT) Inc. Forward or reverse spiked primer oligonucleotides targeting individual viruses were synthesized on a 10 nmol scale in 2-8 pools with standard desalting and 6 nmol of each individual oligonucleotide was mixed and then resuspended to a final volume of 500 μl in IDTE pH 8.0 (IDT).

We prepared quantitative internal process controls (iPC) by spiking both commercially available reference material, the ZymoBIOMICS Fecal Reference (Zymo Research, D6323), and an aliquot of one of our wastewater RNA extractions, with varying concentrations of HeLa cell RNA as process controls.

### 5.7 Library preparation

RNA was used for metagenomic sequencing because it captures both RNA viruses and transcriptionally active DNA viruses. Each sample underwent two parallel library preparation workflows for direct comparison: one using standard unbiased metagenomic next-generation sequencing (mNGS) with random hexamers only, and one using MSSPE with both spiked primers and random hexamers. Library preparation followed protocols described in previous MSSPE and manufacturer protocols of the NEBNext Ultra II Library Prep Kit (New England Biolabs E7771S), with the following additions: External RNA Controls Consortium (ERCC) RNA Spike-In Mix at 25 pg/µL (Invitrogen, Waltham, MA) as internal quantitative controls, FastSelect -5S/16S/23S (Qiagen) at 1:10X ratio for bacterial ribosomal RNA depletion, and random hexamers. Finally, 1.75 µL of sample RNA was added to 3.25 µL of the master mix for fragmentation and priming.

The master mix for fragmentation and priming for MSSPE libraries, was spiked with targeted primers in addition to random hexamers in a 1:2 volume ratio . Dry lyophilized oPools (Integrated DNA Technologies) were resuspended to achieve a final stock of 100 µM. Sequencing libraries were constructed from the purified cDNA using the NEBNext Ultra II Library Prep Kit.

### 5.8 Metagenomic sequencing

After library prep, we performed quality control measures to assess the amount and integrity of input nucleic acid prior to sequencing using a Qubit 4 Fluorometer (Thermo Fisher), and performed electrophoresis visualization on a 1.8% agarose gel.

Libraries were pooled using a two-step normalization process adapted from Mayday et al^52^. Briefly: libraries were first pooled in equal volumes and sequenced on an iSeq 100 at low depth to determine the relative representation of each barcoded library in the total pool. Using the ratio of reads per sample to total reads, a normalization factor was calculated and applied to determine the volume of each library required for equimolar pooling in the final sequencing run. Perfect equimolar representation was not achieved due to inherent variation on insert size between different samples, and limitations of available library volume. The normalized library pool was then sequenced on an Illumina NextSeq 2000 or NovaSeq X 1.5B 1 Lanes (0.8 billion clusters) to generate 150 bp, paired-end sequences.

### 5.9 Microbial identification and bioinformatic analysis

The free open-source cloud-based bioinformatic pipeline and metagenomic analysis platform CZ ID (czid.org) was used to process raw metagenomic sequencing data and classify taxonomy^2^. During data preprocessing, CZ ID removes low quality reads, filters out human reads and randomly subsamples unique QC- and human-filtered reads to 2 million reads (reported abundance values reflect non-deduplicated reads and, thus, may exceed 2 million preprocessed reads per sample). Preprocessed reads are then used for assembly and both reads and contigs are used for taxonomic assignment. Taxonomic assignment is based on searches against NCBI’s nucleotide (NT) and non-redundant (NR) protein databases. CZ ID’s sample report summarizes detected taxa and their abundance based on the number of reads (including those mapped to contigs) aligning to a given taxon.

We applied a mass-normalized background model, as implemented in CZ ID, to account for contamination. The background model is used to calculate the relative abundance of taxa detected in negative controls. The relative abundance is normalized by an estimated input mass obtained from the relationship between known ERCC concentrations and sequencing reads. The relative abundance is then used to calculate a z-score metric to evaluate if a given taxon is above background levels in samples relative to negative controls. We used z-score > 1 and average alignment length > 50 base pairs thresholds for assigning taxa. Additionally, detected taxa needed to be supported by at least 1 reads per million (rPM), 3 reads, or 1 contig aligning to NT and/or NR databases. Taxa that did not meet these thresholds were removed from the analysis.

Unique reads aligning to targeted viral taxa were exported from CZ ID and uploaded to Geneious Prime (v2025.1.3) to generate genome coverage statistics. The default Geneious mapper was used at highest sensitivity. Reads from MSSPE libraries were trimmed by 15 nucleotides at the 5′ and 3′ ends to remove any artefacts of spiked primers before mapping to the most closely matched reference genomes. Coverage statistics were exported from Geneious Prime and genome coverage was calculated in R, where the number of reference bases covered were divided by the total reference genome length. Genome coverage of PBoV species mapped to PBoV 1, accession no. KY489985. Genome coverage of PCV species mapped to PCV2, accession no. OL677619. Genome coverage of EV-G species mapped to accession no. MZ328119. Genome coverage of HPV18 mapped to HPV18 E6/E7 genes, accession no. M20325. Genome coverage of PKV mapped to accession no. NC_027054. Genome coverage of PAstV species mapped to Porcine astrovirus 4 strain, accession no. NC_023675. Genome coverage of PSapeV species mapped to Sapelovirus A, accession no. ON375367. Genome coverage of PTeV species mapped to Teschovirus A, accession no. OL944705.

### 5.10 Statistical analysis

Reads per million (rPM) were calculated by dividing the number of reads aligning to a taxon in the NCBI NT or NR database by the total number of reads minus the number of ERCC reads per sample, and multiplying this result by one million. The ERCC-normalized rPM was used for all downstream analyses. For each paired sample, enrichment was quantified as the ratio of rPM obtained from the MSSPE library relative to the corresponding mNGS library. Non-detected targets were assigned a small pseudocount rPM = 0.1 for log10 fold-change calculation. This value is conservative and well below the lowest non-zero rPM observed, ensuring negligible influence on detected taxa. The pseudocount permits stable ratio-based comparisons for targets detected by only one method, without introducing directional bias because it was applied symmetrically across samples and library preparations. Results were robust to alternative small pseudocount values. Enrichment was calculated as:

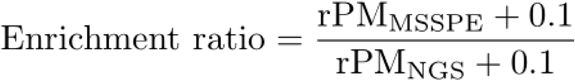

Median, mean, and interquartile range (IQR) fold change values are reported. Because enrichment ratios span several orders of magnitude, we presented enrichment boxplots using a log10-scaled y-axis. Percent increase in genome coverage was calculated as the absolute difference between coverage obtained with MSSPE and random hexamers (mNGS) alone. Wilcoxon signed-rank test assessed paired differences for fold change in rPM across samples as well as absolute difference in genome coverage between library prep methods across viral targets, with P < 0.05 considered statistically significant. Multiple-testing correction (Benjamini–Hochberg) was applied to virus-level comparisons only. We calculated the rank-biserial correlation (r) as an effect size measure from the paired Wilcoxon signed-rank test. Effect sizes provide a sample-size–independent measure of enrichment magnitude and consistency, which can be more informative for evaluating library preparation performance in paired technical replicate designs. We used McNemar’s test on paired samples for each virus and calculated odds ratios based on discordant detections, which were then pooled across all viruses to obtain an overall odds ratio with 95% confidence intervals.

Taxon-level metagenomic read counts were downsampled to target sequencing depths using random subsampling without replacement. For each sequencing depth, 15 independent rarefactions were generated per sample and divided by library preparation method. The alpha-diversity metric observed richness was computed for each replicate to assess diversity recovery across sequencing depths. The Friedman rank test was used to determine statistical differences among sequencing depths. Viral target detection was evaluated by calculating the proportion of rarefaction replicates where read counts met or exceeded defined detection thresholds (5, 10, 15 reads). To assess whether library preparation method influenced virus detection probability across different sequencing depths, we used a generalized linear mixed-effects model with a binomial distribution. The model can be written as:

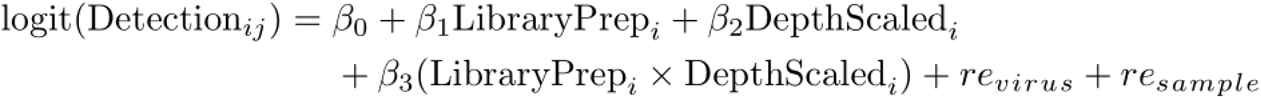

where *Detection* is the binary outcome, *i* is a specific sample, *j* is a specific viral target, *LibraryPrep* is the effect of MSSPE vs mNGS, *DepthScaled* is the effect of sequencing depth scaled by 100,000 reads, *LibraryPrep* x *DepthScaled* is the interaction between library prep and sequencing depth, *re_virus_* is a random effect for viral species, and *re_sample_* is a random effect for samples. Random intercepts for virus target and sample were included to account for differences in baseline detectability among viruses and non-independence of replicate measurements within samples. Model overdispersion was evaluated using a Pearson residual–based statistic. Variance explained was quantified using Nakagawa’s marginal and conditional R² values. Data processing and visualization were implemented in R (v.4.5.0), using multiple packages (ggplot2, tidyverse, vegan, phyoseq, patchwork, kableExtra, purrr, lme4, ggpubr, stringr, performance, viridis, superheat).

## 6. DATA AVAILABILITY

Data supporting the findings of this study are included in the manuscript. Any additional information required to reanalyze the data reported in this work is available from co-corresponding authors. Sequence data have been deposited in the NCBI Sequence Read Archive (NCBI BioProject accession no. PRJNA1391017). MSSPE primer sequences tested in this study are provided in Supplementary Data 1. Protocols are available at https://docs.google.com/document/d/1q0rzj-1f2XUMUGpyVHeTR5srC9z0A7dolWU20n_7Fow/

## 7. CODE AVAILABILITY

Unobfuscated primer design algorithm code *Open MSSPE Design* is available for download and unrestricted usage on GitHub: https://github.com/opendream/open-msspe-design Code used in data analysis has been deposited on GitHub: https://github.com/mattparkerls/metagenomic-spiked-primer-enrichment-paper

## ACKNOWLEDGEMENTS

This study was supported by pilot funds from the Skoll Foundation. We acknowledge the generous support and mentorship from the Rapid Response group at the Chan Zuckerberg Biohub San Francisco, with the donation of reagents, including synthesized primers, and sequencing services used for this research.

## 8. AUTHOR CONTRIBUTIONS

M.C.P. conceived, designed, and supervised the study. M.C.P., T.Y., P.S., and T.E. collected samples. N.A., K.S., and A.M. extracted samples and prepared libraries. J.G. performed QC, sequencing, advised wet lab protocols, edited manuscript and helped with everything. K.C. mentored, advised bioinformatic analysis, and edited manuscript. M.C.P., S.U., and P.P. designed and built the bioinformatic pipelines including Open MSSPE Design software. M.C.P. performed metagenomic analyses. S.H.O. helped with editing manuscript and offering helpful suggestions. M.S. helped with project initiation and mentorship. N.G.D. helped with experimental design, manuscript reviews, and mentorship. M.C.P. and P.S. helped with funding acquisition. M.C.P. wrote the manuscript draft. All authors read, reviewed, and approved the manuscript.

## SUPPLEMENTARY TABLES

**Supplementary Table 1.** Sequencing metrics for all non-control samples.

| Sample | Reads (mNGS) | Reads (MSSPE) | Passed Filter % (mNGS) | Passed Filter % (MSSPE) | DCR (mNGS) | DCR (MSSPE) | Viral Reads % (mNGS) | Viral Reads (MSSPE) | Non-phage Reads % (mNGS) | Non-phage Reads % (MSSPE) |
| --- | --- | --- | --- | --- | --- | --- | --- | --- | --- | --- |
| s11 | 51,727,246 | 50,818,782 | 6.79 | 6.28 | 1.76 | 1.59 | 12.20 | 18.90 | 2.16 | 3.50 |
| s17 | 31,579,706 | 47,462,476 | 27.86 | 10.15 | 4.41 | 2.42 | 14.64 | 15.93 | 3.23 | 5.21 |
| s15 | 38,795,268 | 93,010,162 | 18.81 | 11.12 | 3.64 | 5.17 | 11.09 | 14.06 | 2.29 | 4.69 |
| s7 | 518,666 | 1,273,898 | 40.78 | 47.84 | 1.31 | 1.32 | 15.39 | 13.22 | 5.96 | 2.87 |
| s12 | 55,612,598 | 45,556,186 | 7.34 | 8.51 | 2.03 | 1.94 | 9.22 | 10.53 | 1.19 | 1.66 |
| s8 | 1,199,608 | 4,303,916 | 59.53 | 63.22 | 1.41 | 1.49 | 8.99 | 8.95 | 1.69 | 2.62 |
| s14 | 45,011,752 | 61,293,528 | 7.64 | 7.36 | 1.72 | 2.25 | 6.74 | 6.96 | 0.79 | 1.25 |
| s13 | 43,297,796 | 46,082,456 | 10.33 | 9.82 | 2.23 | 2.28 | 7.51 | 6.95 | 0.95 | 1.21 |
| s19 | 81,398,780 | 75,310,306 | 3.59 | 3.74 | 1.46 | 1.41 | 3.83 | 6.81 | 2.12 | 1.90 |
| s20 | 24,564,082 | 79,608,770 | 15.66 | 4.01 | 1.92 | 1.60 | 1.52 | 6.20 | 1.15 | 3.19 |
| s16 | 31,110,126 | 54,777,360 | 7.94 | 4.70 | 1.24 | 1.29 | 1.62 | 5.76 | 0.33 | 1.22 |
| s6 | 10,097,840 | 7,722,606 | 35.73 | 38.83 | 1.79 | 1.50 | 2.31 | 4.97 | 0.07 | 0.13 |
| s2 | 455,222 | 2,221,908 | 50.17 | 82.17 | 1.33 | 1.43 | 3.90 | 4.76 | 0.78 | 0.66 |
| s4 | 1,001,116 | 1,204,728 | 66.06 | 84.58 | 1.38 | 1.36 | 4.19 | 4.51 | 2.22 | 2.53 |
| s18 | 56,664,844 | 55,901,566 | 6.81 | 5.01 | 1.93 | 1.40 | 1.62 | 3.13 | 0.52 | 0.82 |
| s5 | 8,270,204 | 23,493,784 | 53.27 | 33.34 | 1.66 | 1.96 | 2.07 | 2.55 | 0.09 | 0.12 |
| s9 | 24,845,878 | 3,502,792 | 16.82 | 81.15 | 2.10 | 1.49 | 0.70 | 1.42 | 0.41 | 0.67 |
| s10 | 8,850,592 | 54,021,306 | 46.24 | 10.33 | 2.06 | 2.77 | 0.36 | 0.53 | 0.12 | 0.28 |

<sup>a</sup>The duplicate compression ratio (DCR) is calculated as the total number of reads after QC filtering and host/human read removal divided by the number of unique reads.
<sup>b</sup>Viral reads includes phages.
<sup>c</sup>Non-phage reads includes all viral taxa with phages removed.

**Supplementary Table 2.** Comparison of microbial diversity between library preparation and sample type

| **Diversity Measure** | **NGS** | **MSSPE** | **p.value** | **Type** |
| --- | --- | --- | --- | --- |
| Shannon | 5.50 | 5.73 | 0.01 | Slurry |
| Simpson | 0.98 | 0.98 | 0.09 | Slurry |
| Richness | 5,216.14 | 6,319.36 | 0.02 | Slurry |
| Shannon | 5.65 | 6.09 | 0.12 | Wastewater |
| Simpson | 0.98 | 0.98 | 0.12 | Wastewater |
| Richness | 8,573.00 | 8,832.88 | 0.62 | Wastewater |
| Shannon | 5.53 | 5.81 | 0.00 | Overall |
| Simpson | 0.98 | 0.98 | 0.01 | Overall |
| Richness | 5,962.11 | 6,877.92 | 0.01 | Overall |

**Supplementary Table 3.**
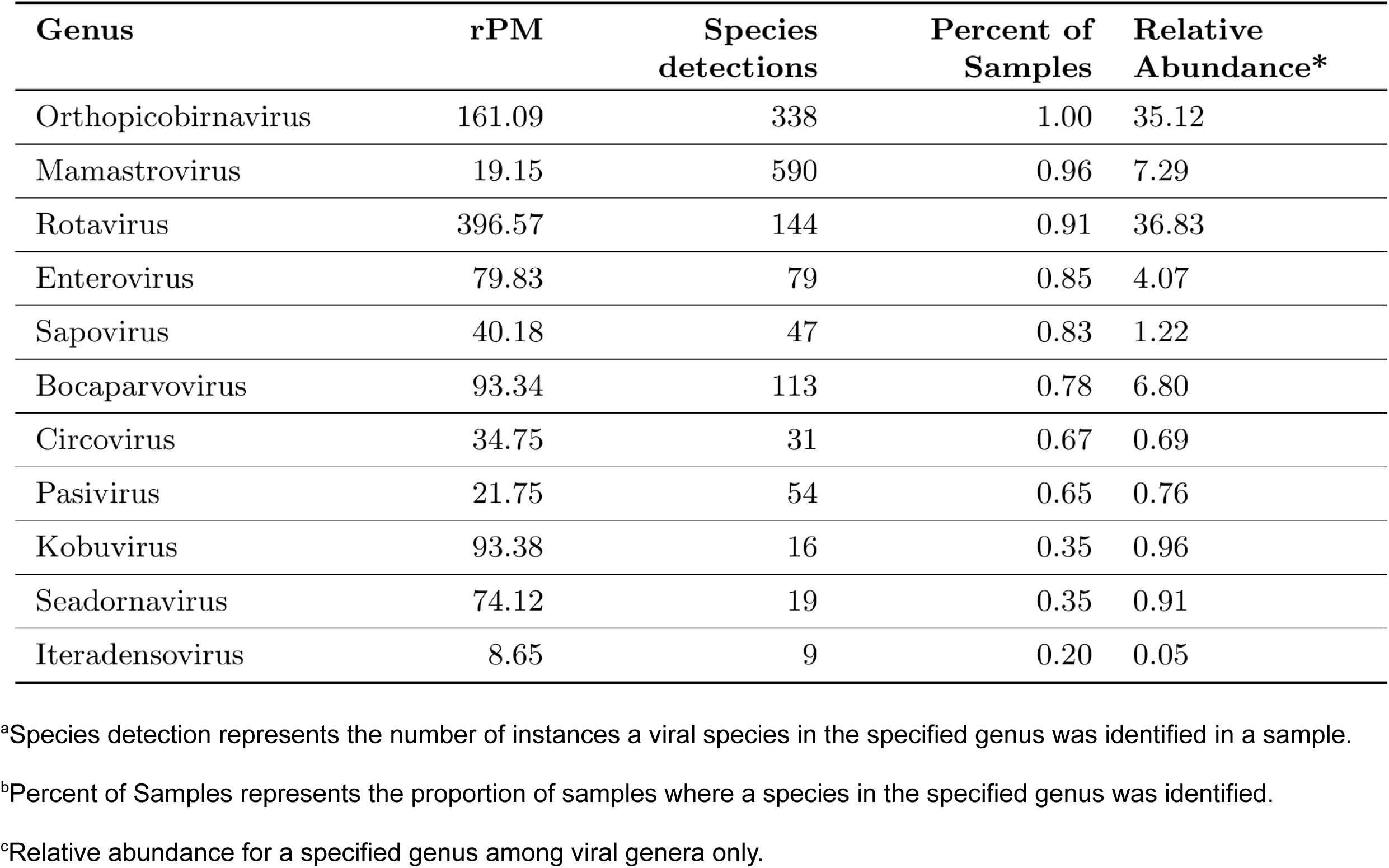
Most abundant viral genera detected across all samples.

| **Genus** | **rPM** | **Species**<br>**detections** | **Percent of**<br>**Samples** | **Relative**<br>**Abundance\*** |
| --- | --- | --- | --- | --- |
| Orthopicobirnavirus | 161.09 | 338 | 1.00 | 35.12 |
| Mamastrovirus | 19.15 | 590 | 0.96 | 7.29 |
| Rotavirus | 396.57 | 144 | 0.91 | 36.83 |
| Enterovirus | 79.83 | 79 | 0.85 | 4.07 |
| Sapovirus | 40.18 | 47 | 0.83 | 1.22 |
| Bocaparvovirus | 93.34 | 113 | 0.78 | 6.80 |
| Circovirus | 34.75 | 31 | 0.67 | 0.69 |
| Pasivirus | 21.75 | 54 | 0.65 | 0.76 |
| Kobuvirus | 93.38 | 16 | 0.35 | 0.96 |
| Seadornavirus | 74.12 | 19 | 0.35 | 0.91 |
| Iteradensovirus | 8.65 | 9 | 0.20 | 0.05 |

<sup>a</sup>Species detection represents the number of instances a viral species in the specified genus was identified in a sample.
<sup>b</sup>Percent of Samples represents the proportion of samples where a species in the specified genus was identified.
<sup>c</sup>Relative abundance for a specified genus among viral genera only.

**Supplementary Table 4.** Proportion of rarefaction replicates where read accounts met or exceeded 5 reads aligned to the target viral taxa.

| Virus | Library<br>Prep | 0.1M | 0.3M | 0.5M | 0.7M | 0.9M | 1.1M | 1.3M | 1.5M |
| --- | --- | --- | --- | --- | --- | --- | --- | --- | --- |
| EVG | MSSPE | 0.38 | 0.77 | 0.86 | 0.92 | 0.92 | 0.92 | 0.92 | 0.92 |
| EVG | NGS | 0.28 | 0.70 | 0.77 | 0.83 | 0.88 | 0.87 | 0.90 | 0.92 |
| PAstV | MSSPE | 1.00 | 1.00 | 1.00 | 1.00 | 1.00 | 1.00 | 1.00 | 1.00 |
| PAstV | NGS | 1.00 | 1.00 | 1.00 | 1.00 | 1.00 | 1.00 | 1.00 | 1.00 |
| PBoV | MSSPE | 1.00 | 1.00 | 1.00 | 1.00 | 1.00 | 1.00 | 1.00 | 1.00 |
| PBoV | NGS | 0.87 | 0.96 | 1.00 | 1.00 | 1.00 | 1.00 | 1.00 | 1.00 |
| PCV | MSSPE | 0.54 | 0.83 | 0.92 | 0.92 | 0.92 | 0.92 | 0.92 | 0.92 |
| PCV | NGS | 0.32 | 0.61 | 0.67 | 0.71 | 0.72 | 0.74 | 0.75 | 0.75 |
| PKV | MSSPE | 0.00 | 0.08 | 0.17 | 0.24 | 0.25 | 0.25 | 0.25 | 0.25 |
| PKV | NGS | 0.00 | 0.00 | 0.00 | 0.01 | 0.01 | 0.03 | 0.08 | 0.08 |
| PSapeV | MSSPE | 0.57 | 0.58 | 0.58 | 0.58 | 0.58 | 0.58 | 0.58 | 0.58 |
| PSapeV | NGS | 0.48 | 0.62 | 0.72 | 0.72 | 0.75 | 0.75 | 0.75 | 0.75 |
| PSapoV | MSSPE | 0.67 | 0.75 | 0.75 | 0.76 | 0.76 | 0.78 | 0.80 | 0.80 |
| PSapoV | NGS | 0.53 | 0.90 | 0.97 | 0.99 | 1.00 | 1.00 | 1.00 | 1.00 |
| PTeV | MSSPE | 0.49 | 0.67 | 0.67 | 0.67 | 0.67 | 0.67 | 0.67 | 0.67 |
| PTeV | NGS | 0.38 | 0.53 | 0.59 | 0.62 | 0.65 | 0.66 | 0.67 | 0.67 |

<sup>a</sup>Detection probability was then summarized as a function of sequencing depth and library preparation method.

**Supplementary Table 5.** Fixed and random effects from the generalized linear mixed-effects model of species richness.

| Effect | Estimate | SE | z | p | OR | CI<br>25% | CI<br>75% | Percent<br>change |
| --- | --- | --- | --- | --- | --- | --- | --- | --- |
| Library<br>prep | 0.182 | 0.020 | 8.946 | 0.000 | 1.200 | 1.153 | 1.249 | 19.990 |
| Sequencing<br>depth | 0.032 | 0.002 | 19.762 | 0.000 | 1.032 | 1.029 | 1.035 | 3.209 |
| Sample<br>type | -0.609 | 0.170 | -3.575 | 0.000 | 0.544 | 0.390 | 0.760 | -45.587 |
| Lib<br>prep *<br>Se-<br>quenc-<br>ing<br>depth | -0.012 | 0.002 | -5.300 | 0.000 | 0.988 | 0.984 | 0.992 | -1.188 |
| Lib<br>prep *<br>Sample<br>type | -0.040 | 0.023 | -1.760 | 0.078 | 0.961 | 0.919 | 1.005 | -3.935 |
| Sequencing<br>depth *<br>Sample<br>type | 0.015 | 0.002 | 8.160 | 0.000 | 1.015 | 1.011 | 1.018 | 1.472 |
| Lib<br>Prep *<br>Depth *<br>Sample<br>Type | 0.004 | 0.003 | 1.759 | 0.079 | 1.004 | 0.999 | 1.009 | 0.445 |

<sup>a</sup>Fixed effects show the estimated influence of library preparation method, sequencing depth (scaled per 100,000 reads), sample type, and their interaction on species richness.
<sup>b</sup>Estimates are presented on the log-odds scale, with corresponding standard errors, z-values, p-values, odds ratios (OR = $\exp(\beta)$ ), and 95% confidence intervals for the OR.

## Supplementary Figures

**Supplementary Figure 1.**
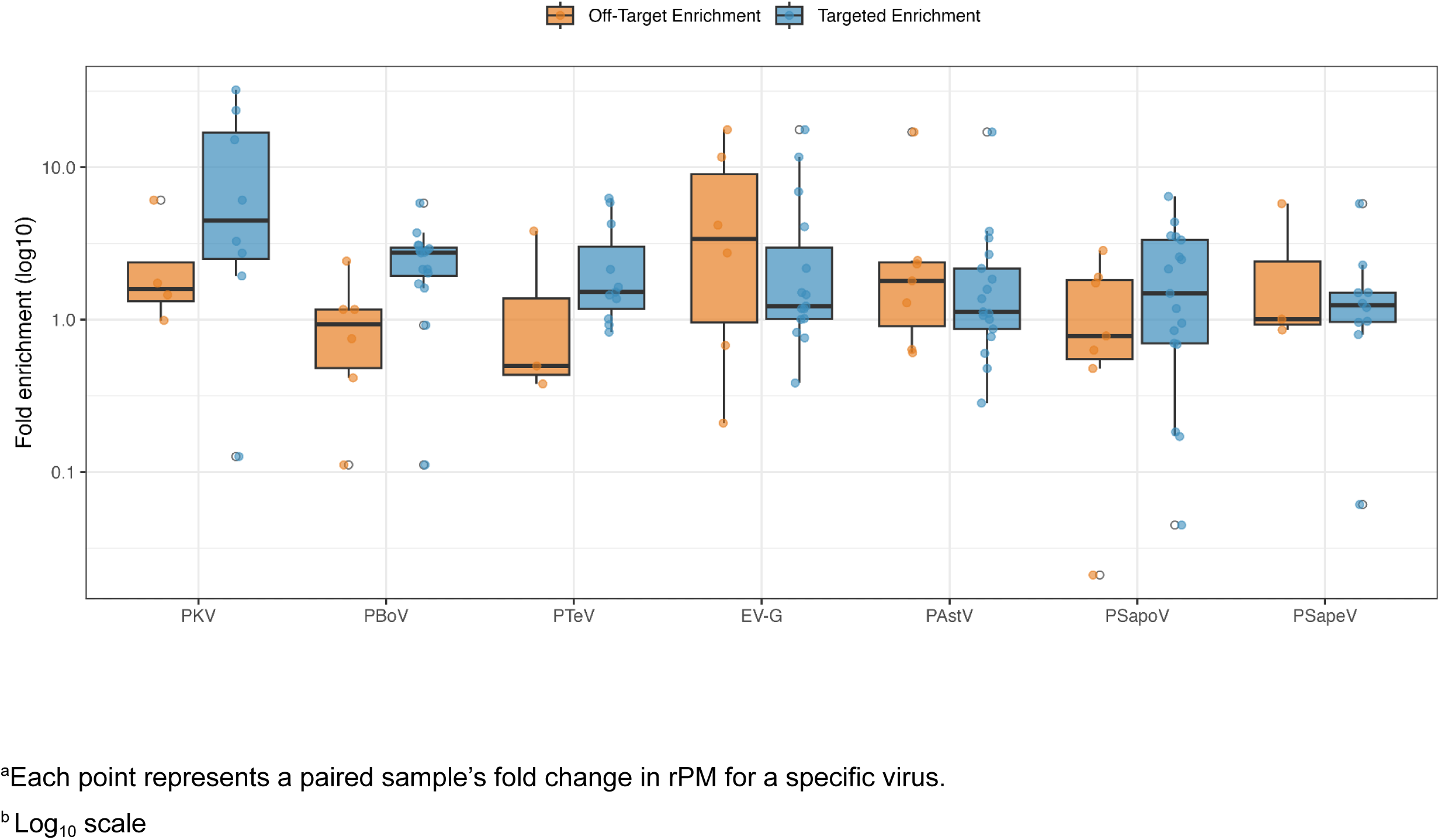
Fold change in median reads per million (rPM) aligned to target viruses with random hexamers and non-target spiked primers (orange) compared to random hexamers and target spiked primers (blue).

**Supplementary Figure 2.**
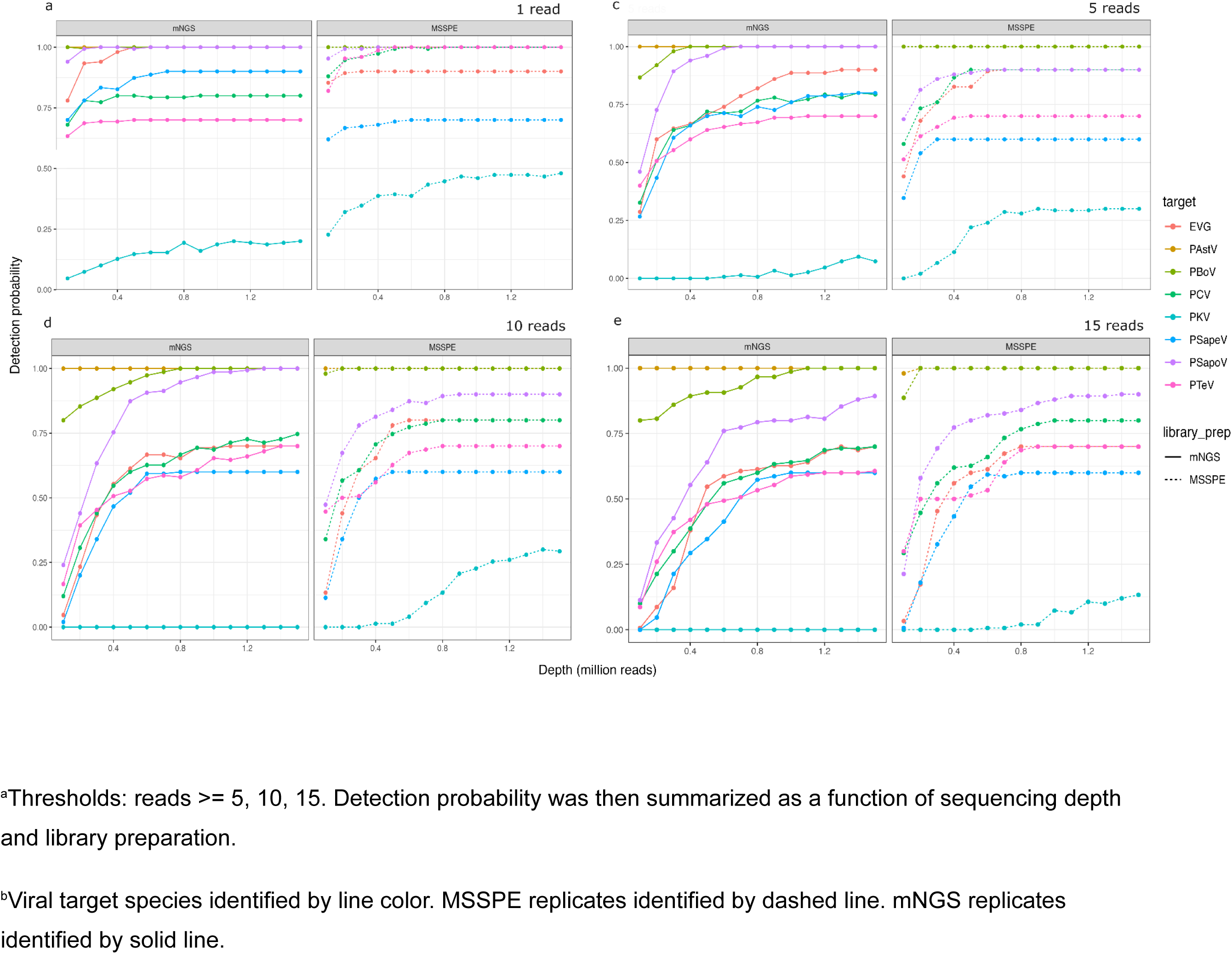
Proportion of rarefaction replicates where viral taxa met or exceeded defined detection thresholds.

**Supplementary Figure 3.**
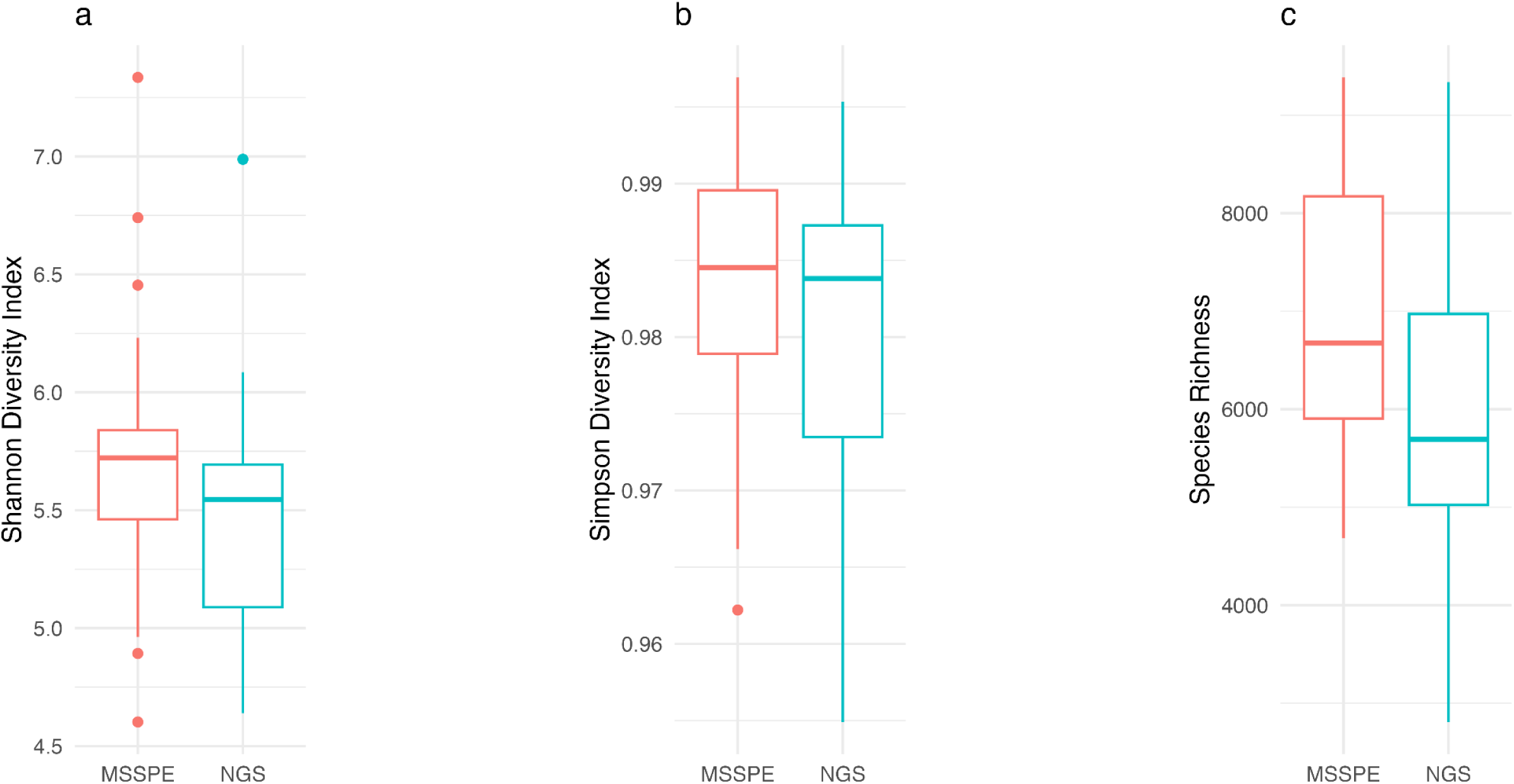
Overall species diversity index values for all phyla across all samples grouped by library preparation method according to Shannon diversity, Simpson’s diversity, and Species richness.

